# Unveiling the unique interaction mechanism of herpes simplex virus 2 glycoprotein C with C3b

**DOI:** 10.1101/2025.10.04.680448

**Authors:** Moisés Hasim Rojas Rechy, Doina Atanasiu, Lauren M Hook, Tina M Cairns, Wan Ting Saw, Adam Cahill, Zilin Guo, Antonio N. Calabrese, Neil A Ranson, Harvey M Friedman, Gary H Cohen, Juan Fontana

**Affiliations:** School of Molecular and Cellular Biology, Faculty of Biological Sciences and Astbury Centre for Structural and Molecular Biology, University of Leeds, Leeds, United Kingdom; Instituto Biofisika, CSIC-UPV/EHU, Leioa, Bizkaia, Spain; School of Dental Medicine, University of Pennsylvania, Philadelphia, USA; Perelman School of Medicine, University of Pennsylvania, Philadelphia, USA

**Author notes:** MHRR and DA contributed equally to this work. Corresponding authors: GHC JF.

## Abstract

The complement cascade is part of the first line of defence against viral infections, and many viruses have evolved to block it. For example, glycoprotein C (gC) from Herpes Simplex Virus 1 and 2 (gC1 and gC2) facilitates infection by modulating the complement cascade through an interaction with C3b. gC is also involved in attachment and other viral processes. However, our understanding of the molecular mechanisms of gC have been limited due to the absence of a structure. AlphaFold predicts that gC contains a disordered N-terminus and three immunoglobulin-like domains. Here, we generated various gC2 constructs and demonstrated that gC2 domains 1 and 2 are necessary and sufficient to interact with C3b and block the alternative pathway. A gC2 construct lacking the N-terminus in complex with C3b was characterised by cryo-EM at 3.6 Å, providing the first structure for gC2, and revealing that the interaction is predominantly driven by gC2 domain 2 and the MG8 domain of C3b. This structure was confirmed by cross-linking mass spectrometry and by using C3b-blocking antibodies that recognised gC2 linear epitopes at the interface with C3b. Overall, the gC-C3b interaction is different from other C3b-interacting partners, providing a novel mechanism to regulate the complement cascade.

## Main

During infection, the host immune response targets the incoming pathogen. Specifically, the complement cascade is part of the innate immune system and involves many plasma proteins that recognise pathogens and induce inflammatory responses to fight infection^1,2^. Within these, C3 and its derivatives play a central part in amplifying the immune response^3^. As pathogens, many viruses have evolved the ability to block the complement cascade to facilitate infection^4^. For example, glycoprotein C (gC) from herpes simplex virus (HSV) was the first viral protein identified to interact with the complement protein C3b^5,6^; however, understanding its mechanism of action has been hindered by the lack of a structure. Additionally gC mediates viral attachment, by interacting with heparan sulfate^7^; and is involved in antibody blocking, protecting HSV entry glycoproteins from neutralising antibodies^8–10^. Thus, gC is an important protein for HSV infection.

Herpes simplex virus 1 and 2 (HSV-1 and HSV-2) are dsDNA viruses that belong to the *Herpesvirales* order. While HSV infection is normally associated with mild blisters or ulcers, HSV-1 can lead to severe meningitis and encephalitis^11^ and HSV-2 infection is associated with an increase in the risk of acquiring and transmitting HIV^12^. Given that there are currently no approved vaccines for HSV and that treatments only alleviate the symptoms, but cannot eradicate infection, further research on the molecular biology of HSV infection is required to tackle the burden caused by these viruses, which affect over 64% (HSV-1) and 13% (HSV-2) of the world-wide population^13^.

To evade the complement cascade, glycoprotein C (gC) from both HSV-1 and HSV-2 (termed gC1 and gC2, respectively), interact with C3b and other C3-related proteins^14^. Strikingly, biochemical studies indicate that while both gCs bind to C3b, the biological outcomes of this binding differ: gC1 accelerates the decay of the C3 convertase by inhibiting properdin binding^15,16^, whereas gC2 stabilizes the convertase^14^. In any case, both gC1 and gC2 provide protection against complement-mediated neutralization of viral infectivity^17^. For example, it has been shown that a gC-null HSV-1 is 100-fold less virulent than WT *in vivo*^18^, and that gC1 is involved in evading antibody-independent and antibody-dependent complement-mediated virus inactivation and lysis of virus-infected cells^19,20^.

Different studies have shed light on our understanding of the structure/function of gC. For example, the use of gC truncations defined two main functional domains on gC: an N-terminal domain that for gC1 (but not for gC2) blocks C5 and properdin binding to C3b^21^ and a C3b-binding domain^21–23^, which based on competition studies has been suggested to bind to a similar location on C3b for both gC1 and gC2^23^. Furthermore, amino acid insertion experiments have determined the C3b binding regions for gC1 and gC2^22,23^. However, all these studies were hampered by the lack of a structure for gC. Such a structure would allow us to understand how gC interacts with C3b and provide novel ways to block HSV infection and regulate complement activation. Furthermore, a gC structure would allow the rationalization of a gC2-based mRNA vaccine, which together with gD2 and gE2 has been shown to provide protection against both HSV-1 and HSV-2 infection^24^.

Herein, we used AlphaFold^25^ to predict the structures of gC1 and gC2 bound to C3b, which suggested that both gCs adopt a similar structure and bind to C3b using a similar mechanism of interaction. Based on the AlphaFold-predicted domains, we generated different gC2 constructs, which were tested for binding to C3b and for blocking the alternative and classical pathways of the complement cascade. These results suggest that only domains 1 and 2 of gC2 are required for C3b binding. Functional *in vitro* assays showed that the same gC2 constructs that bind C3b block the alternative pathway at a stage prior to the C5 convertase activity, suggesting the gC2 binding to C3b is sufficient for complement inhibition. Finally, cryo-electron microscopy (cryo-EM) and crosslinking mass spectrometry (XL-MS) were employed to structurally characterise the gC2-C3b complex, and this structure was confirmed using gC2 antibodies that block binding to C3b. This structure suggests that gC binds to C3b using a novel mechanism that could be exploited therapeutically.

### AlphaFold provides a plausible model for the gC-C3b interaction

gC1 and gC2 have been determined to have similar C3b binding regions^22,23^ and to bind to a similar location on C3b^23^. Furthermore, their sequences are 68% identical (Extended Data Fig. 1), suggesting that they share a similar structure. To improve our understanding of how gC interacts with C3b, we generated AlphaFold^25^ models of gC1 and gC2 ectodomains in complex with C3b (Fig. 1A and Extended Data Fig. 2). These resulted in interface predicted template modelling (ipTM) scores of 0.69 and 0.79 for gC1 and gC2, respectively (predictions with values above 0.8 are considered to be of high-quality; values between 0.6 and 0.8 require further validation). AlphaFold suggests that gC contains a long, disordered N-terminus (N-t) (which lowers the ipTM scores) and 3 immunoglobulin-like domains, which we termed domains 1 to 3, from N-t to C-terminus (Fig. 1, Extended Data Fig. 1).

**Figure 1.**
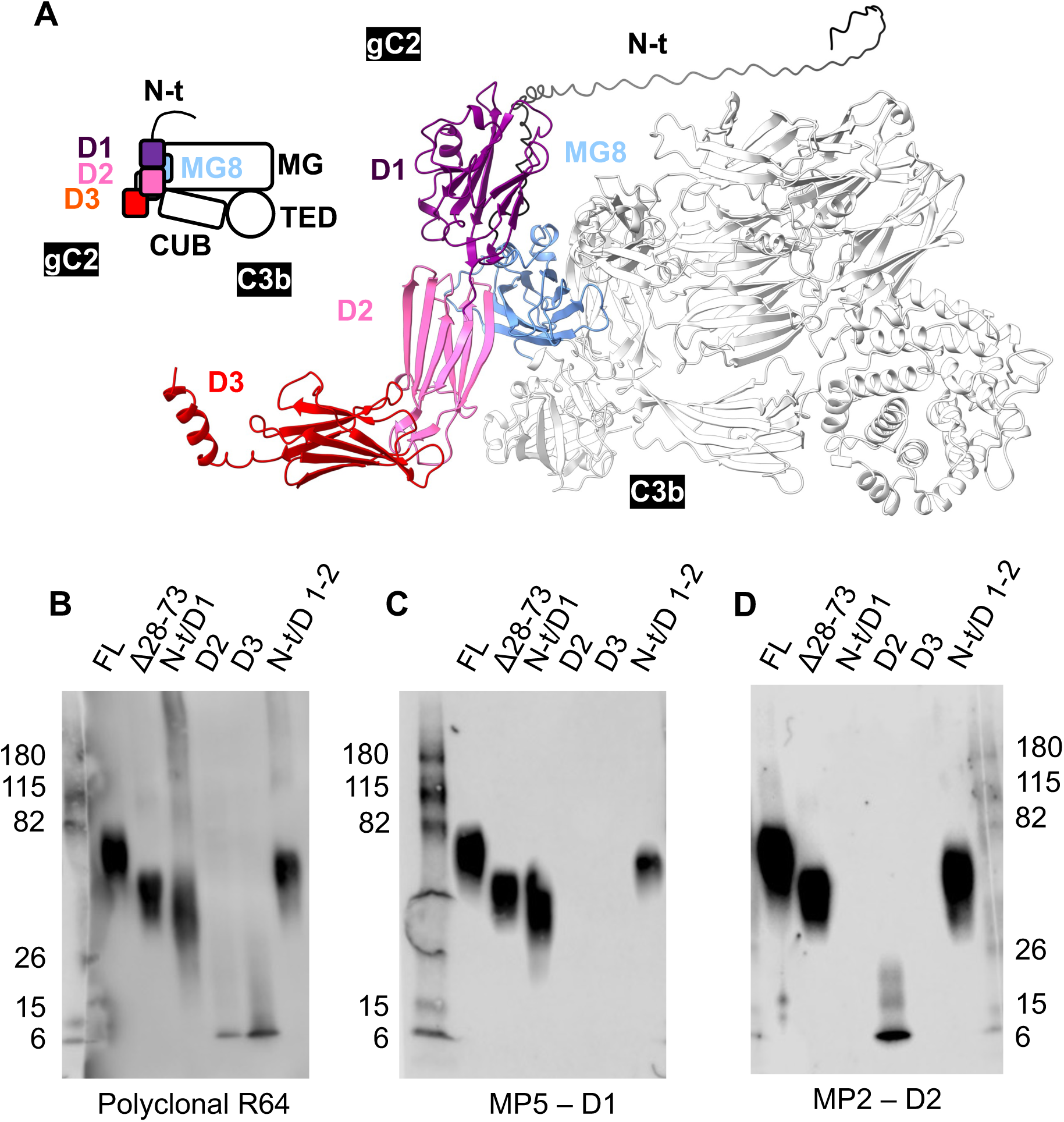
AlphaFold prediction of the gC2-C3b complex and expression of gC2 constructs. A) AlphaFold gC2-C3b complex prediction shown as a cartoon (left) and as ribbons (right). gC2 domains are represented as follows: N-t, black; D1, purple; D2, pink; D3, orange. C3b is in white, except for the MG8 domain, which is blue. B-D) Native SDS-PAGE and western blot to test the correct folding of the gC2 constructs. B) Polyclonal R64. C) Conformational MP5 antibody, which recognises D1. D) Conformational MP2 antibody, which recognises D2.

Notably, the AlphaFold model fits well with the previously described C3b binding regions of gC, which were based on amino acid insertions affecting C3b binding. For gC2^23^ three regions responsible for binding were previously identified. Region I (amino acids 102 to 107) is within D1, close to MG8 in the AlphaFold prediction; region II (222-279) is at the core of D1 and D2, so this region might be required to maintain the proper folding of these domains; and region III (307-379) includes the D2 beta strand that is proposed to interact with C3b and part of D3 (Extended Data Fig. 1). The identified gC1 C3b binding regions are similar to those from gC2^22^: region I corresponds to 124-137 (93-106 on gC2), region II to 276-292 (245-261 on gC2) and region III to 339-366 (308-335 on gC2). However, an additional region was identified, region IV, corresponding to residues 223-246 (192-215 on gC2); this region is located between regions I and II, at the core of D1 in the AlphaFold prediction (Extended Data Fig. 1).

Overall, these results suggest that both gC1 and gC2 fold and bind to C3b in a similar way, as it has been suggested in the literature (e.g.^17,22^).

### gC2 truncations based on the AlphaFold prediction fold correctly

For further experiments we decided to focus on gC2 since: 1) the AlphaFold prediction of gC2-C3b interaction has better metrics than the gC1-C3b prediction; 2) gC2 has a 10-fold higher affinity for C3b compared to gC1^26^; 3) the AlphaFold predictions suggest that both gC1 and gC2 fold and bind to C3b in a similar way; and 4) gC2 is currently one of the immunogens in a HSV trivalent vaccine that is in human trials^27^.

To test the AlphaFold prediction, we designed six different gC2 ectodomain constructs that were expressed and purified (Extended Data Fig. 3). Full-length gC2 ectodomain was expressed as a control (termed gC2-FL). A version of gC2 that lacked most of the N-terminus (gC2 Δ28-73) was expressed to test if this region is involved in the interaction with C3b. Domains 1 (including the N-terminus), 2 and 3 were expressed independently, to test their individual contributions to C3b binding (N-t/D1, D2 and D3). Given the AlphaFold prediction suggests that only domains 1-2 make direct contact with C3b, we also expressed a version of gC2 containing the N-terminus and these two domains (N-t/D1-2). For all the constructs, a His-tag was added for purification purposes.

Next, to validate if the constructs were correctly folded, we used conformational antibodies on a native SDS-PAGE and western blot (to keep the constructs structured)^28^. As expected, all the constructs were detected by the gC2-specific polyclonal antibody R64^14^, further confirming that they are correctly expressed (Fig. 1B). Additionally, conformational monoclonal antibodies MP5 and MP2 have been described to bind to regions corresponding to D1 and D2, respectively^29^. Indeed, all gC2 constructs containing D1 (i.e. gC2 FL, Δ28-73, N-t/D1 and N-t/D1-D2) were detected by MP5, suggesting they are correctly folded (Fig. 1C). Similarly, constructs containing D2 (i.e. gC2 FL, Δ28-73, D2 and N-t/D1-D2) were detected by MP2, also suggesting that they are correctly folded (Fig. 1D). Of note, we could not assess the folding of D3, because there is no antibody available that maps to this domain.

To further validate the proper folding of the gC2 constructs, we took advantage of circular dichroism (CD), which allows the identification of the secondary structure of proteins, given that alpha-helices, beta-strands and disordered regions result in characteristic CD spectra. Specifically, β-sheets are characterised by having a low peak at around 220 nm, while disordered regions have negative peaks at around 195 nm^30^. When performing CD on gC2 FL, its spectra had a low peak at around 210 nm, which can be explained based on the long-disordered N-t combined with the β-sheet-rich D1-3 (Extended Data Fig. 4A). Compared to gC2 FL, the spectra of Δ28-73 is shifted to the right (i.e. fewer disordered regions), given this construct lacks the disordered N-t, while the spectra of N-t/D1-D2 is shifted to the left (i.e. more disordered regions), given that the ratio of disordered regions to β-sheet is increased due to the lack of D3 (Extended Data Fig. 4A). Similarly, the spectra of gC N-t/D1 is shifted to the left compared to those of D2 and D3, given the disordered to β-sheet ratio is increased in this construct compared to D2 and D3 (Extended Data Fig. 4B).

### gC2 D1 and D2 are required for C3b binding

Once we had confirmed that the gC2 constructs fold correctly, we tested which ones could bind C3b. To this end we used an enzyme-linked immunosorbent assay (ELISA). C3b was immobilized in 96-well plates, and individual gC2 constructs were incubated separately. Binding of gC2 constructs to C3b was detected using the gC2-specific polyclonal antibody R64, followed by a secondary antibody and a colorimetric reaction (Extended Data Fig. 4C). The results showed that gC2 FL can bind C3b, as expected (Fig. 2B). Additionally, the gC2 N-t/D1-2 and Δ28-73 constructs can also bind C3b. On the other hand, isolated individual domains from gC2 (N-t/D1, D2 and D3) did not bind C3b, (Fig. 2B).

**Figure 2.**
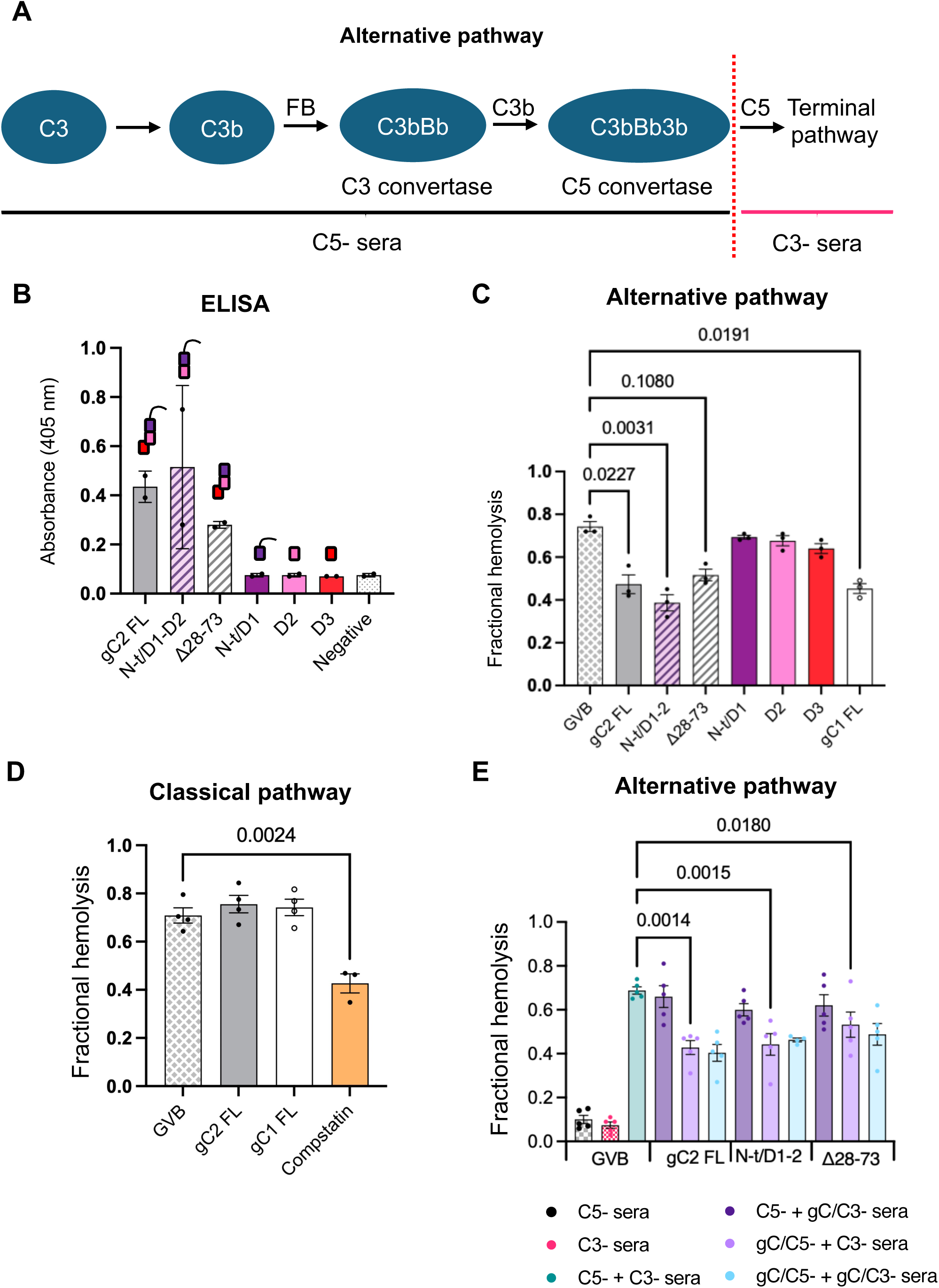
Functional characterisation of the gC2 constructs. A) Schematic of the alternative pathway. B) ELISA to determine C3b binding of the gC2 constructs. Data are from 2 biological repeats (n=2), each measured in triplicate (3 technical repeats). C) Assay to determine inhibition of the alternative pathway by the different gC constructs. GVB was used as a positive control for haemolysis. Data are from 3 biological repeats (n=3). D) Assay to determine inhibition of the classical pathway by the different gC constructs. GVB was used as a positive for haemolysis and compstatin as a known inhibitor. Data are from 4 biological repeats (n=4), except for compstatin, which are from 3 biological repeats (n=3). E) Assay to determine if inhibition of the alternative pathway by the gC constructs occurs prior to C5 convertase activity or after. Data are from 5 biological repeats (n=5), except for N-t/D1-2 C5-/gC+C3-/gC which is from 4 biological repeats (n=4). For all panels, data points are presented as mean values and bars represent standard deviation.

To confirm these results, a biosensor experiment was performed. gC2 constructs that bound to C3b (gC2 FL, Δ28-73 and N-t/D1-2) and a negative control (N-t/D1) were immobilized onto a His chip (CM5) independently, taking advantage of their His-tags. Then, C3b was injected, and binding was detected as an increase in the response units (RUs) (Extended Data Fig. 4D). These results confirmed the ELISA data, since gC FL, N-t/D1-2 and Δ28-73 bound to C3b, while individual domains did not (Extended Data Fig. 4E).

Overall, these results show that both gC2 D1 and D2 are required for C3b binding, while the N-t and D3 are dispensable. This fits well with the literature, since a gC1 construct lacking the N-terminus (up to residue 123) and D3 (residues 367 to 469) has been shown to bind to C3b^15^.

### gC2 binding to C3b results in inhibition of the complement alternative pathway

gC modulates complement activation mainly via the alternative pathway^16^ (Fig. 2A). Therefore, to test the regulatory properties of our gC2 constructs, we performed an assay to evaluate the activity of the alternative complement pathway (Fig. 2C). In this assay, rabbit erythrocytes are incubated with human serum, resulting in activation of the alternative complement pathway and haemolysis. However, incubation in the presence of inhibitors of the alternative pathway reduces the extent of haemolysis. In the absence of any alternative pathway inhibitors, the assay resulted in ∼75% haemolysis. However, when this assay was performed in the presence gC1 FL and gC2 FL, haemolysis was reduced to ∼50%, confirming the role of both gC1 and gC2 in inhibiting the alternative pathway. Additionally, gC2 N-t/D1-2 and Δ28-73 reduced haemolysis in a comparable level to gC2 FL, while N-t/D1, D2 and D3 did not inhibit haemolysis (Fig. 2C). These results suggest that gC2 binding to C3b, as determined above, is sufficient for inhibition of the alternative pathway. As expected, gC1 FL and gC2 FL did not reduce haemolysis in a modified CH50 assay, which is used to assess inhibition of the complement classical pathway by measuring the extent of haemolysis of sensitised sheep erythrocytes. However, compstatin, a drug known to inhibit the classical pathway^31^, did (Fig. 2D).

Next, to pinpoint the step at which gC2 exerts its inhibition, we used a modified AP50 assay^32^. Here, rabbit erythrocytes are first incubated with C5-depleted sera, allowing for assembly of the C3 convertase, activity of the C3 convertase and assembly of the C5 convertase at the surface of erythrocytes, while C5 convertase activity and progression of the cascade is prevented due to the absence of C5 as a substrate. After washing, erythrocytes are incubated with C3-depleted serum, providing C5 as a substrate for the C5 convertase and therefore haemolysis, while preventing the formation of new C5 convertases due to the absence of additional C3/C3b. Inhibitors can be present at any of these two steps, allowing us to discern if their effect is up to the formation of the C5 convertase or if they regulate the cascade from the C5 convertase activity. When only C5-or C3-depleted sera were used in this assay, no hemolysis could be detected, since each sera by itself lacks all the components for the alternative pathway activation. However, when erythrocytes were consecutively incubated with the two sera, ∼70% hemolysis occurred (Fig. 2E). When gC2 FL was present in both C5- and C3-depleted sera, a haemolysis reduction similar to that seen in the standard AP50 assay (to ∼45%) was observed. A similar reduction was observed when gC2 FL was present only in the C5-depleted sera, which was not observed when gC2 FL was only present in the C3-depleted sera, suggesting gC2 blocks the C3 convertase assembly, the C3 convertase activity, or C5 convertase assembly. Similar results were observed for gC2 N-t/D1-2 and Δ28-73, suggesting all constructs have a similar mechanism of action, and that the main complement cascade blocking activity of gC resides within D1 and D2.

### gC2 D1 and D2 interact with the MG8 domain from C3b while gC2 N-t interacts with MG3

To solve the structure of gC2-C3b by cryo-EM we initially attempted to characterise gC2(426t), a truncated version of gC2 extensively used for biochemical assays^33^, mixed with C3b in a 2:1 ratio (Extended Data Fig. 5). While this resulted in some 2D classes showing extra density compatible with being gC at the site suggested by AlphaFold (Extended Data Fig. 5B, top right), downstream image processing resulted in an average containing only C3b (Extended Data Fig. 5C), likely due to only a small fraction of C3b molecules interacting with gC2 in this sample and preferred orientation issues.

Therefore, to stabilise the complex, we took advantage of the crosslinker BS3^34^. Additionally, we decided to use gC2 Δ28-73, given that: 1) gC2 Δ28-73 binds to C3b and blocks the alternative pathway in a manner comparable to gC2 FL (Fig. 2); 2) a gC1 construct lacking the N-t (up to residue 123) inhibits complement activation in vivo^35^, and due to the structural similarities between gC1 and gC2 we reasoned that the same would be true for gC2; 3) the gC N-t is suggested to be flexible by the AlphaFold prediction (Extended Data Fig. 2), making gC2 Δ28-73 more amenable for structural characterisation.

When gC2 Δ28-73 and C3b were mixed in a 2:1 ratio in the absence of crosslinker, bands for gC2 Δ28-73 (∼60 kDa), and for the C3b chains (∼115 kDa for the α-chain and ∼75 kDa for the β-chain, which can run at an apparent lower molecular weight in reducing conditions^36^) were visible by SDS-PAGE (Extended Data Fig. 6A, lane 0). As the crosslinker concentration was increased from a 25 molar excess to 100 molar excess, bands at the expected size of a gC2-C3b complex (∼240 kDa) were observed. This was accompanied by the disappearance of the C3b bands, suggesting all C3b is present as part of the gC2-C3b complex (Extended Data Fig. 6A, lanes 25, 50 and 100). We next sought to confirm the results from SDS-PAGE using mass photometry, which uses a beam of light that interacts with a molecule in solution that can be correlated to the size of the molecule^37^. In the absence of the crosslinker, free gC2 Δ28-73 (∼19% of the sample) and C3b (∼39% of the sample) and a 1:1 complex (∼45% of the complex) were detected (Extended Data Fig. 6B). However, when the crosslinker was added, no free C3b was detected; instead, only free gC2 Δ28-73 (∼38% of the sample) and a 1:1 complex (∼51% of the sample) were detected (Extended Data Fig. 6C). We reasoned that this sample was ideal for cryo-EM, since all C3b molecules would be forming a gC2-C3b complex, and gC2 would not affect particle-picking or downstream image processing due to its small size. Therefore, we proceeded with single-particle cryo-EM imaging of this sample, using 25 molar excess of BS3 to minimise the formation of aggregates.

Cryo-EM grids of gC2 Δ28-73 – C3b crosslinked with BS^3^ were prepared by plunge-freezing, and data acquired. To overcome issues with preferred orientation that we observed in the gC2(426t)-C3b dataset, data was collected at 0° and at 25° tilts^38^. ∼35,000 movies were collected corresponding to ∼470,000 particles, which were processed to generate a ∼3.6 Å map of the whole gC2-C3b complex, and a local refined map of the gC2-C3b interface at an overall resolution of ∼4.0 Å (Extended Data Figs. 7 and 8 and Extended Data Table 1). Of note, the gC2-C3b interactions were resolved at local resolution of ∼3.0-3.5 Å resolution (Extended Data Fig. 7F and G and Extended Fig. 8A and B), allowing us to confidently model this region.

The atomic model of gC2 D2 and of the C3b-contacting regions of D1 confirmed the AlphaFold prediction (Fig. 3 and Extended Data Table 1). Furthermore, analysis of the gC2-C3b interactions by PDB PISA^39^ confirmed that the key interactions are located within amino acids 95-110 of gC2 D1 (corresponding to the previously termed binding region 1^23^) mostly interacting with amino acids 1417-1431 of the MG8 domain from C3b (Extended Data Fig. 9, yellow); and amino acids 322-331 of gC2 D2 (previously termed binding region 3) mostly interacting with amino acids 1449-1462 of C3b MG8 (Extended Data Fig. 9, green). These interactions are driven by hydrogen bonds and salt bridges between both proteins (Fig. 3B to D). Of note, given the intrinsic flexibility of some regions of C3b, namely TED, CUB and C345c^3^, these were barely visible in our cryo-EM maps. Similarly, there was no clear density for gC2 D3, suggesting it is also flexible (Extended Fig. 8A and B). Additionally, extra densities around N331 could be seen, likely due to glycosylation of this residue (Extended Fig. 10).

**Figure 3.**
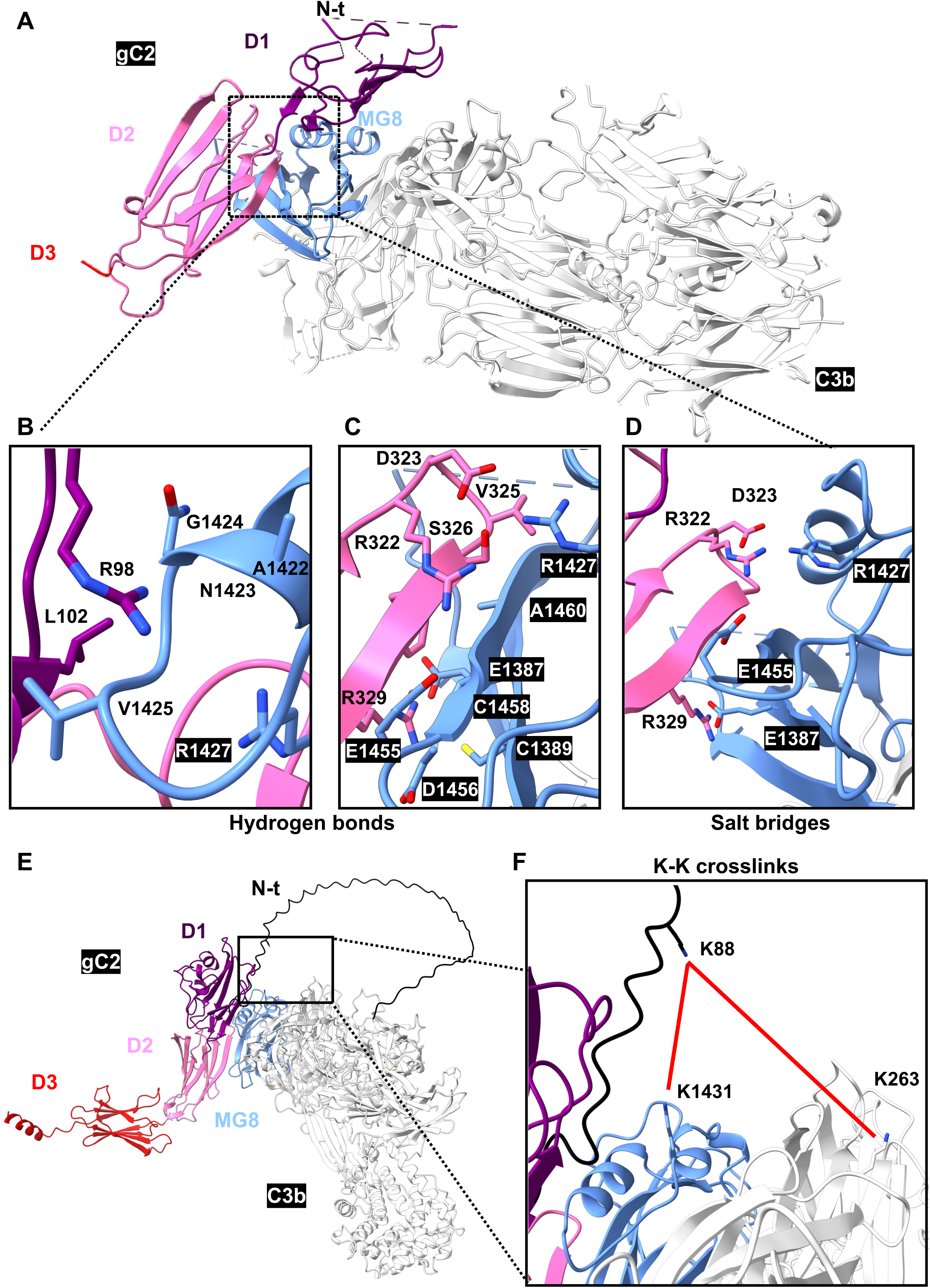
Structure of the gC2 Δ28-73-C3b complex. A) Ribbon representation of the gC2 Δ28-73-C3b complex as determined by cryo-EM. The square highlights the interactions between the two proteins, as determined by PDB-PISA. B and C) Hydrogen bonds (black dashed lines) formed between D1 (B) or D2 (C) and MG8. D) Salt bridges (black dashed lines) formed between D2 and MG8. E and F) XL-MS detected Lys-Lys crosslinks (red lines) overlaid on the gC2-C3b AlphaFold prediction. Domain colouring is as in Figure 1.

As expected, there was also no density in the cryo-EM map for the small N-t region of gC2 Δ28-73, further suggesting that this is a flexible region. To gain an insight of its location and interactions, we took advantage of cross-linking mass spectrometry (XL-MS), which allows the identification of cross-linked peptides between two interacting proteins. When gC2 FL-C3b was crosslinked using DSBU (disuccinimidyl dibutyric urea), which can form lysine-lysine crosslinks that are up to 30 Å apart, we identified 2 crosslinks driven by Lys 88 from gC2 (Fig. 3E and F and Extended Data Fig. 11): one with Lys 1431 from C3b MG8, and the other one with Lys 263 from MG3, close to the interface with MG7 and MG6. These crosslinks suggest potential interactions that place the N-t of gC2 close to the binding region of the nanobody hC3Nb1^40^ and the antibody S77^41^, which are known C3b inhibitors, and suggest a potential mechanism of inhibition for gC2. Additionally, XL-MS also identified an interaction between gC2 Ser 326 (within D2) and C3b Lys 1368 (within MG8) (Extended Fig. 8C and D), further confirming the interactions happening between both domains.

### gC2-C3b interaction can be blocked by gC2 antibodies, confirming the atomic model

To confirm the gC2-C3b atomic model, we scanned our gC antibody library against 15 amino-acid long overlapping gC peptides, aiming to identify antibodies recognising linear gC2 regions. For this, biotinylated peptides were immobilized either on streptavidin coated plates for ELISA or printed on a streptavidin chip for surface plasmon resonance (SPR) screening (using Carterra LSA). By ELISA, antibody H1291 recognized gC2 peptide 321-335 (Extended Data Fig. 12A). By SPR, antibody H1196 recognized a single gC1 peptide (137-151) and, to a lesser extent, the equivalent gC2 peptide 105-119 (Extended Data Fig. 12B and C). This suggests that the H1196 epitope might adopt a more conformation-sensitive presentation in gC2. Mapping of these two antibodies with gC2 truncations confirmed the location of H1291 epitope to D2 and the requirement of both D1+D2 for H1196 recognition (Extended Data Fig. 12D). The location of these two antibodies suggests that they should affect the gC2-C3b interaction (Extended Data Fig. 9C). To measure their effect on this interaction, we used SPR and a Biacore 1k+ instrument (Cytiva) (Extended Data Fig. 12E). gC2 antibodies were captured on a ProG chip, followed by the sequential flow of soluble gC2 FL and C3b. After the initial antibody capture, the binding of gC2 and C3b are registered as two separate increases in response units (RUs) if the antibody does not interfere with C3b binding. However, if the antibody competes, there will be no increase in RUs after the C3b injection. When gC2 was captured by the LH50a antibody (manuscript in preparation), C3b bound, suggesting that it does not interfere with the gC2-C3b complex formation (Fig. 4A). However, when gC2 was captured by the MP2 antibody, which does not block C3b binding (manuscript in preparation) (Fig. 4B), H1291 (Fig. 4C) or H1196 (Fig. 4D), no C3b binding was observed. These results confirm that the gC2 regions between 95-110 and 322-331, as determined in our structure, are key for binding to C3b, validating our atomic model of the complex, and provide potential targets on gC to regulate HSV infection.

**Figure 4.**
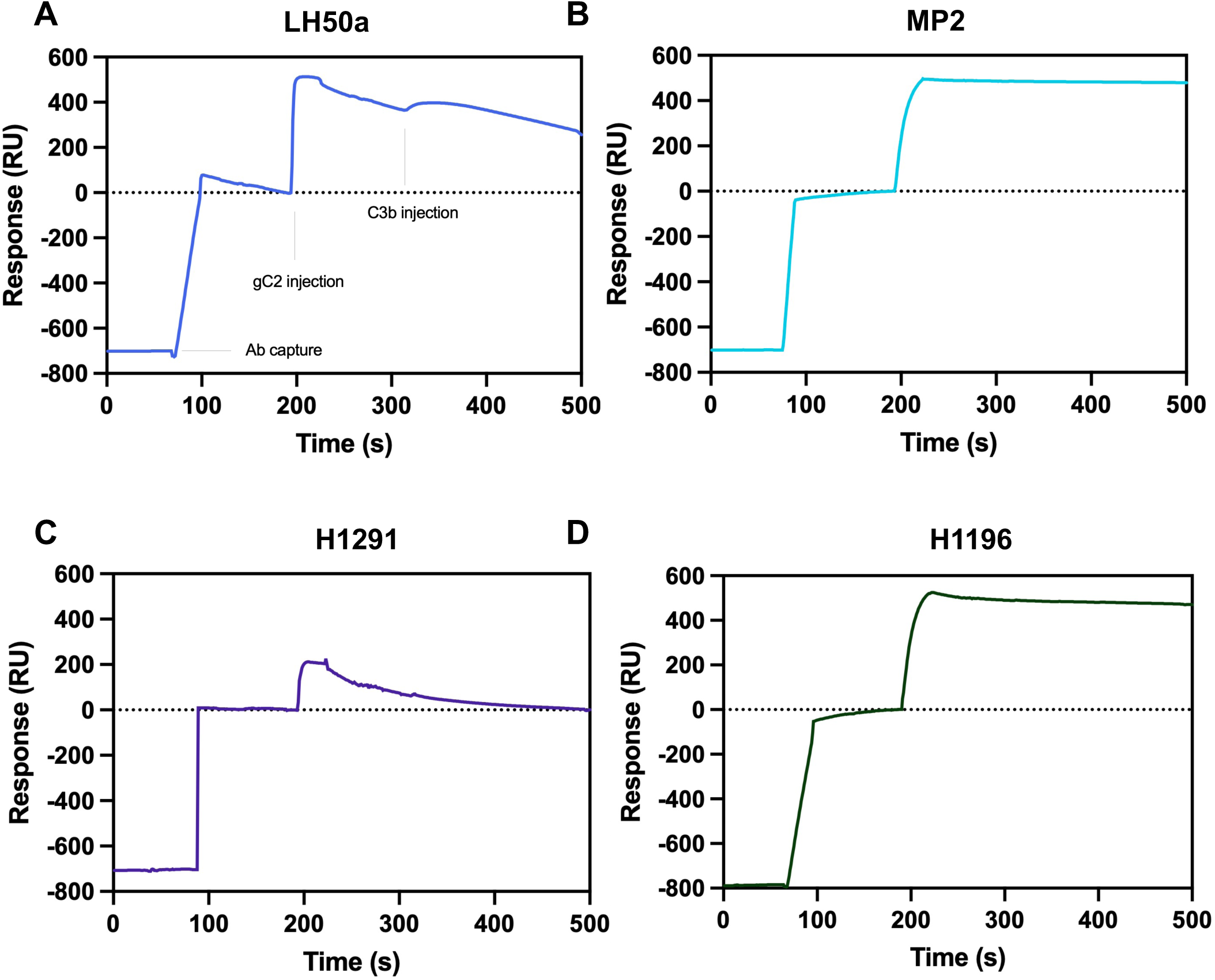
Antibody blocking of the gC2-C3b interaction. SPR results to detect C3b blocking using gC2 antibodies. Antibodies were captured on a ProG chip, followed by the sequential flow of soluble gC2 FL and C3b. Curves show initial antibody binding followed by gC2 binding. No increase in the RU after C3b injection shows blocking. A) LH50a antibody, does not block gC2-C3b interaction (negative control). B), Results for the MP2 antibody, known to block the gC2-C3b interaction (positive control). C) Results for H1291. D) Results for H1196.

## Discussion

The complement system is a key defence mechanism and as such it is not surprising that it is regulated by many viruses. Specifically, HSV gC is known to interact with C3b, blocking the complement cascade. However, our molecular understanding of this process has been limited due to the lack of a gC-C3b structure. Here, we designed different gC2 constructs based on AlphaFold predictions of the gC-C3b complex, and showed that gC2 D1 and D2 are necessary and sufficient for binding to C3b and blocking the alternative pathway. Additionally, we solved the structure of gC2-C3b using cryo-EM, and showed by XL-MS that the gC2 D1-D2 interaction with C3b places the disordered gC2 N-t towards the C3b region were known inhibitors bind.

Several viruses encode molecules that inhibit complement, including HSV-1^6^, HSV-2^26^, Kaposi’s sarcoma-associated herpesvirus (KSHV)^42^ and vaccinia virus^43^. For example, KSHV regulator of complement activation (RCA) homolog (termed Kaposica) resembles factor H, a key complement regulator^42^. Additionally, human cytomegalovirus infection has been shown to increase cell surface expression of host RCAs, also regulating the complement^44^. Strikingly, gC is different to known complement regulators; instead, it is similar to the ectodomains of many cell surface receptors, as determined using FoldSeek^45^. This includes nectin-1, one of the known HSV receptors^46^. Therefore, alphaherpesvirus gC could have its origin in a cellular surface protein, which then evolved D1 and D2 to interact with C3b, resulting in its unique mode of binding. gC’s origin as a receptor also explains its role in attachment to heparan sulfate, which has been shown to locate within amino acids 129-160 of gC1, within D1^47^.

gC is present across the whole alphaherpesvirus subfamily (e.g.^48^). Predictions for different alphaherpesvirus gCs in the Big Fantastic Virus Database^49^ always contain a disordered N-t (of different lengths), which might be involved in sterically prevent antibody interaction with viral glycoproteins^8–10^, and 3 immunoglobulin-like domains. In fact, a similar organisation for different gC proteins was proposed using structure prediction algorithms^50,51^. Overall, these predictions agree with our gC2 structure and point to a conserved structure and function of gC across alphaherpesviruses, and therefore a similar mechanism of interaction with C3b.

Additionally, both gC1 and gC2 have been shown to bind to C3, albeit with lower affinity than to C3b^21^. This also agrees with our gC-C3b binding model, since the MG8 is present in C3, but the side that interacts with gC is occluded by the anaphylatoxin domain^52^, which likely needs to be displaced by gC to form the gC-C3 complex.

While both gC1 and gC2 have been shown to provide protection against complement-mediated neutralization of viral infectivity^17^ and to bind to C3b in a similar way^22^, significant differences in the consequences of this interaction have been described in the literature. gC1 N-t has been shown to block properdin (and C5) binding, therefore destabilising the C3 convertase^21^. On the other hand, gC2 has been described to stabilise the C3 convertase by an unknown mechanism^14^. Additionally, it has been suggested that both proteins might regulate the complement cascade in a different way, given only gC1 has been shown to disrupt activation of alternative pathway C3 convertase, C3bBb^19,53^. Our results using modified CH50 and AP50 assays shed light in this matter, and indicate that gC1 and gC2 similarly inhibit the alternative pathway, rather than the classical pathway. This again aligns with our structural observations (for gC2) and predictions (for gC1), which suggest that both proteins bind C3b in a similar way. Potential differences with previous results are likely due to the assays used. Differences between gC1 and gC2 could also be due to the role of the N-t domain, or to their quaternary structure: while a putative gC1 dimer has been proposed^54^, we found no evidence of gC2 dimers by negative staining or mass photometry. In any case, a putative gC dimer would have to monomerise before interacting with C3b, since the D1 and D2 interfaces that interact with C3b would be occluded in the dimer.

All together, we propose the following mechanism of inhibition. As previously described, and supported by our structural data, gC contains two main regions with respect to complement regulation: the disordered N-t (which for gC1 is termed the C5/properdin inhibitory binding domain in the literature) and the C3b-binding region (D1 and D2)^21^. Taking into account the differences in sequence and function between gC1 and gC2 N-t regions, we suggest that the N-t-C3b interactions, and therefore their potential complement regulatory activities, are gC-dependent. In any case, we show that the C3b-binding region is necessary and sufficient to bind C3b and block the alternative pathway at a stage prior to C5 convertase activity. Given that the gC D1-D2 binds away from the Factor Bb and properdin binding sites and the C3 convertase catalytic site^55^, we suggest that the gC C3b-binding region by itself does not block or inhibit C3 (pro-)convertase assembly or activity. Based on this, we propose that gC blocks assembly of the C5 convertase, for which a structure is currently unavailable. Therefore, therapeutics blocking the C5 convertase assembly could be developed based on gC D1-D2 and used as complement cascade inhibitors, paving the way for a new type of complement therapeutics.

## Methods

### gC sequences and gC2 constructs

gC1 and gC2 sequences are based on Uniprot entries Q8UYE2 and P06475, respectively. gC2 constructs had the following coordinates: FL, 28-445, which corresponds to the ectodomain after removing the signal peptide (residues 1-27) and up to the transmembrane helix (starting at 448); Δ28-73, 74-445; N-t/D1, 28-236; D2, 237-342; D3, 336-445; and N-t/D1-D2, 28-342. The corresponding DNA sequences were synthesized and cloned by GenScript into pcDNA3. For purification purposes a 6× His tag was added at the C-term of all constructs. Proteins were expressed and purified by GenScript from supernatants of CHO transfected cells using Nickel columns.

### gC Alphafold modelling and metrics

The AlphaFold3 server^25^ was used to model gC1-C3b and gC2-C3b. The C3b models fit well with published C3b structures (e.g. pdb 2I07^56^), with RMSD of ∼1.1 and ∼1.2 Å for gC1-C3b and gC2-C3b, respectively. In these models, gC contains a long flexible N-terminus, as suggested by the predicted local distance difference test (pLDDT), which is a per-residue measure of local confidence (values above 70 correspond to confident predictions, while values above 90 are considered very high confidence); and does not interact with C3b, based on the predicted aligned error (PAE), which is a measure of the confidence in the relative position of two residues within the predicted structure model (the smaller the error, the higher the confidence two domains interact with each other). Specifically, both N-ts are associated with very low pLDDT (< 50; Extended Data Fig. 2A and B) and very high PAE (> 30 Å; Extended Data. Fig. 2C and D). The rest of gC is arranged as 3 immunoglobulin-like domains. The 3 domains from both gC1 and gC2 are associated with high pIDDT values (over 70, with the core of gC2 D2 presenting values over 90), suggesting a confident prediction, with gC D1 and D2 presenting low PAE values compared to C3b, suggesting D1 and D2 drive the gC2-C3b interaction (Extended Data Fig. 2C and D).

### Western blotting of gC2 constructs

100 ng of purified proteins were run on Novex 10% Tris-Glycine gel under “native” conditions^28^. After transfer, nitrocellulose membranes were probed with 1 µg/ml purified antibodies followed by goat anti-mouse or anti-rabbit peroxidase secondary antibodies, as indicated and developed using Pierce substrate.

### Circular dichroism of gC2 constructs

gC2 constructs were diluted PBS (137 mM NaCl, 2.7 mM KCl, 10 mM Na_2_PO_4_, 1.8 mM K_2_PO_4_, pH 7.4) at a concentration of 0.2 mg/ml. Proteins were loaded into a 1.0 mm cuvette. Measurements were performed and recorded at a range of 180-260 nm wavelength at room temperature using a Chirascan Plus CD spectrometer (Applied Photophysics).

### ELISA

To determine gC2-C3b interaction, Maxisorb plate (Nunc) plates were coated with 10 µg/ml C3b, diluted in PBS overnight at 4°C and then blocked with 5% milk-0.05% Tween-20-PBS (PBS/T) for 1 hour at room temperature. Plates were washed once with PBS and incubated with 10 µg/ml gC2 constructs for 1 hour. Bound gC was detected with anti-gC2 polyclonal antibody R64 (PMID:8053154) followed by anti-rabbit-HRP secondary antibody. For Ab mapping studies, ELISA plates were coated with 10 µg/ml gC2 truncations. Each set of proteins was probed with the indicated Mabs and goat anti-mouse HRP secondary Ab. Detection was performed at 405 nm using a BioTek Synergy H1 plate reader (Agilent Technologies).

### gC2-C3b interaction determined by Biacore

Binding experiments were perfomed using a Biacore 1K+ (Cytivia) at 25°C. Filtered 1x HBS-EP+ (10 mM of HEPES, 150 mM NaCl, 3 mM EDTA, pH 7.4) was used in all experiments. The gC2 constructs were coupled to a CM5-anti His chip (Cytivia) through their C-terminal His-tags using a 10 µg/min flow rate. Different concentrations were used to obtain similar resonance units (RU) as follows: gC2 FL, 66 nM; gC2 Δ28-73, 83 nM; gC2 N-t/D1, 111 nM; gC2 D2, 714 nM; gC2 D3, 6 µM; gC2 N-t/D1-D2, 167 nM. The purified C3b (CompTech) and/or antibodies were injected at a concentration of 1333 nM. Glycine 100 mM pH 2 was used for regeneration to baseline levels.

For antibody blocking experiments, 1 µg/ml of the indicated Mabs were immobilized on a ProG chip (Cytiva) using a Biacore 1k+ instrument (Cytiva). First, gC2 FL was injected at 400 nM, followed by a second injection of C3b at 278 nM. Glycine pH 2 was used to remove Ab-protein complexes from the chip surface until the response signal returned to baseline.

### Alternative pathway assay

Serum from a HSV-1 and HSV-2 seronegative human was diluted with GVBo buffer (0.1 % gelatin, 5 mM Veronal, 145 mM NaCl, 0.025 % NaN3, pH 7.3) to a C3 concentration of 5 µM and mixed with an equal volume of test protein construct at 10 µM in a microtiter plate (final volume of 50 µl). GVBo served as a buffer control. The serum:protein mix was then supplemented with Mg^2+^/EGTA to support activation of the alternative complement pathway (and inactivation of the classical complement pathway). 2.5×10^7^ rabbit erythrocytes were then added to each well for a final well volume of 100 µl. Control wells included blanks (100 µl GVBo alone), cell blanks (cells with GVBo supplemented with Mg^2+^/EDTA) and 100% lysis wells (cells mixed with water). The microtiter plate was incubated for 30 minutes at 37 °C with intermittent mixing. Ice cold PBS (or saline) was added to all wells except the 100% lysis wells. The 100% lysis wells were treated with ddH_2_O. The plate was then centrifuged 400xg, 5 minutes and supernatant was transferred to fresh wells of a microtiter plate. OD was read at 405 nm. Fractional hemolysis was calculated as the OD sample/AVE OD of the 100% lysis wells (cells mixed with water). Normality was determined by the Shapiro-Wilk test. P values were determined by Kruskal-Wallis test with Dunn’s correction for multiple comparisons.

The adapted AP50 assay was similar as above, but took place in two stages. Initially, diluted C5-depleted human serum (Complement Technologies) was mixed with the indicated protein construct, supplemented with Mg^2+^/EGTA to support activation of the alternative complement pathway and then incubated with 2.5×10^7^ rabbit erythrocytes for 30 minutes at 37 °C with intermittent mixing (final well volume of 100 µl; C3 concentration of 1.25 µM/protein construct concentration of 5 µM). Cells were then washed 2 times with GVB supplemented with Mg^2+^/EGTA. Rabbit erythrocytes were resuspended in diluted C3-depleted human serum (Complement Technologies) supplemented with Mg^2+^/EGTA and the indicated protein constructs and then incubated 30 minutes at 37 °C. Ice cold PBS (or saline) was added to all wells except the 100% lysis wells, which were treated with ddH2O. The plates were then centrifuged 400xg, 5 minutes and supernatant were transferred to fresh wells of a microtiter plate. OD was read at 405 nm. Fractional hemolysis was calculated as the OD sample/AVE OD of the 100% lysis wells (cells mixed with water). Normality was determined by the Shapiro-Wilk test. P values were determined by One-way ANOVA with Dunn’s correction for multiple comparisons.

### Classical pathway assay

Serum from a HSV-1 and HSV-2 seronegative human was diluted with GVB++ buffer without azide (0.1 % gelatin, 5 mM veronal, 145 mM NaCl, 0.025 % NaN_3_, 0.15 mM CaCl_2_ and 0.5 mM MgCl, pH 7.3) to a C3 concentration of 0.63 µM and then mixed with an equal volume of test protein construct at 10 µM in a microtiter plate (final volume of 50 µl). GVB++ served as a buffer control. 2.5×10^7^ antibody-sensitized sheep erythrocytes (EA) were then added to each well for a final well volume of 100 µl. Control wells included blanks (100 µl GVB++ alone), cell blanks (cells with GVB++) and 100% lysis wells (cells mixed with water). The microtiter plate was incubated for 30 minutes at 37 °C with intermittent mixing. Ice cold PBS (or saline) was added to all wells except the 100% lysis wells. The 100% lysis wells were treated with ddH_2_O. The plate was then centrifuged 400xg, 5 minutes and supernatant was transferred to fresh wells of a microtiter plate. OD was read at 405 nm. Fractional hemolysis was calculated as the OD sample/OD 100% lysis wells (cells mixed with water). Normality was determined by the Shapiro-Wilk test. P values were determined by unpaired t test.

### Crosslinking gC2 Δ28-73-C3b complex using BS^3^

For cryo-EM purposes, gC2 Δ28-73-C3b and C3b were incubated in a 2:1 molar ratio in PBS at a 62 µM total protein concentration. BS^3^ was dissolved in water to a final concentration of 25, 50 and 100 mM. The crosslinker was added to a 20 µL gC2 Δ28-73-C3b protein complex at room temperature for 30 minutes. The reaction was quenched using 0.1 M of TRIS at room temperature for 15 minutes.

For mass photometry measurements, 4 µL of sample (with and without crosslinker) were diluted into 16 µL of PBS for a final protein concentration of 400 nM. Microscope slides were washed with milli-Q water and dried with a continuous flux of air. Silicone gaskets (CultureWell) were used to place the samples on top of the slides. The measurements were performed using a Refeyn OneMP mass photometer (Refeyn). Measurements were recorded for 60 seconds and the software Discover^MP^ was used to analyse data. A calibration curve was used with standard proteins (66-669 kDa) diluted in 25 mM HEPES pH 7.5, 10 mM NaCl.

### Cryo-electron microscopy grid preparation

To prepare gC2(426t)-C3b and gC2 Δ28-73-C3b grids, 3 µL of sample were applied to Quantifoil 200 Mesh Cu R1.2/1.3 or Quantifoil 300 Mesh Cu R1.2/1.3 graphene oxide grids respectively, previously glow discharged for 30 seconds (PELCO easyGlow). After, sample was vitrified by plunge freezing into liquid ethane using a FEI Vitrobot Mark IV (ThermoFisher). The chamber temperature was set up at 4 °C, 100% humidity, wait time of 1 second and blot force of 1.

### Cryo-electron microscopy data collection

Single particle cryo-EM was performed using a Titan Krios TEM operating at 300 keV and equipped with a Falcon 4 Direct Electron Detector (Thermo Fisher Scientific). Screening and data collection were done using the automated data collection software EPU (ThermoFisher Scientific) gC2 Δ28-73-C3b datasets were collected at 0° and 25° tilts; gC2 gC2(426t)-C3b datasets were only collected at 0°. Movies were collected in electron counting mode, over a defocus range of -0.9 to -2.7 µm and at a nominal magnification of 96,000k, which corresponded to a calibrated pixel size of 0.74 Å/pix.

### Image processing

For both datasets, pre-processing was done using RELION v4.0.0^57,58^, beam-induced motion correction was performed using MotionCor2^59^ and the contrast transfer function (CTF) was calculated using CTTFFIND4^60^. Particles were picked using crYOLO 1.8.0^61^ using the standard model with a threshold of 0.1.

For gC2(426t)-C3b, 7,470 movies were collected. 420,120 particles were initially picked and processed in RELION. Particles were subjected to multiple 2D classification jobs to discard junk particles, resulting in 90,640 “good” particles. The selected particles were used to generate 3 different 3D initial models. The map with the larger number of particles was picked for further processing. 3D classification and auto-refine were used to refine the 90,640 particles.

For gC2 Δ28-73-C3b, 10,087 movies were collected at 0° stage tilt and 14,913 at 25° stage tilt. Datasets were pre-processed independently as above until picking ∼1.4M particles. Further processing was done using CryoSPARC^62^. 2D classification was then performed to remove junk particles, and 886,581 particles were selected and extracted using a 256 px box size (bin 4). For ab initio modelling, 3 classes were used. One of the averages was and used for further 3D classification and the average with the highest number of particles was used for homogeneous refinement. Particles were re-extracted using a 400 px box size (unbinned) and non uniform refinement was performed obtaining a 3.5 Å map. A local refinement job was performed applying a mask for the gC2 area, resulting in a 4 Å map. Post-processing was performed using DeepEMhancer^63^ (used for visualisation purposes only), and local resolution was estimated in CryoSPARC and visualized in ChimeraX^64^.

### Model building and refinement

Initial refinement was performed using molecular dynamics flexible fitting of the AlphaFold atomic model into the cryo-EM map using ISOLDE v1.0^65^. Cycles of manual inspection in Coot^66^ and real-space refinement using Phenix^67^ were then performed until satisfactory results were observed. Model validation was performed using Phenix and MolProbity^68^.

### Crosslinking-mass spectrometry

The gC2 FL-C3b complex was formed at a 1:1 ratio with 4 µM total protein concentration in PBS. DSBU (disuccinimidyl dibutyric urea) was dissolved in DMSO (dimethyl sulfoxide) to a concentration of 20 mM. The crosslinker was added to 20 µL of the gC2 FL-C3b protein complex (final concentration of 1 mM) and the sample was incubated at room temperature for 1 hour.

Cross-linked proteins were then digested using S-Trap micro columns (Protifi), according to the manufacturer’s instructions using MS-grade solvents. First, 25 µL of 10 % (w/v) SDS solution was added to 25 µL of crosslinked protein (concentration of 20 µM). DTT solution (5 µL, 220 mM) was added, and the sample was heated to 95°C for 15 mins and then allowed to cool to room temperature. Iodoacetamide (5.5 µL, 440 mM) was added, and the sample was incubated in the dark for 30 min at room temperature. The solution was acidified by adding phosphoric acid to a final concentration of 1.2 % (v/v). Samples were then diluted with S-Trap binding buffer 7:1 (100 mM triethylammonium bicarbonate (TEAB) pH 7.1 in methanol). Trypsin (Promega) was added from stocks at a concentration of 0.02 µg/µL to a ratio of 1:20 (enzyme:protein). The protein solution containing the appropriate enzyme was loaded onto the S-Trap column and centrifuged at 4000 x g for 30 s, to enable protein capture within the submicron pores of the S-Trap. The column was washed by adding 130 µL of binding buffer before being centrifuged at 4000 x g for 30 s. 30 µL of trypsin (0.02 µg/ µL in TEAB) was added to the top of the S-trap column and samples were incubated at 47 °C for 90 mins. 40 µL of 50 mM TEAB was added and the column was centrifuged at 4000 x g for 1 min. Digested peptides were eluted by sequential washing with of 0.2 % (v/v) formic acid (FA) and then 50 % (v/v) acetonitrile (ACN), with centrifugation (4000 x g for 30 s) after each addition. The eluates from these steps were combined and dried using a Concentrator Plus vacuum centrifuge (Eppendorf).

Dried samples were resuspended in 0.1 % (v/v) trifluoracetic acid to a concentration of ca. 1 µM. Samples were then analysed on a Vanquish Neo LC System (Thermo Scientific) coupled to an Orbitrap Eclipse Tribrid mass spectrometer (Thermo Scientific). The instrument was fitted with an EASY-spray reversed phase LC system with column specifications as follows: particle size: 2 µm, diameter: 75 µm, length: 500 mm, running at 250 nL/min flow and kept at 45°C. Analytes were eluted by a 97 min, 0-50% acetonitrile gradient. The mass spectrometric settings for MS1 scans were: resolution of 120 000, automatic gain control (AGC) target of 3×10^6^, maximum injection time of 50 ms, scanning from 380– 1450 m/z in profile mode. Cycle time was adjusted to 3 seconds and charge states z = 3–8 were isolated using a 1.4 m/z window and fragmented by HCD using optimized stepped normalized collision energies, 30 ±6 volts. Fragment ion scans were acquired at a resolution of 60 000, automatic gain control target of 1×10^5^ ions, maximum injection time of 120 ms, scanning from 200–2000 m/z, underfill ratio set to 1%. Dynamic exclusion was enabled for 30 s (including isotopes). Thermo RAW files were converted to mgf files using Proteome Discoverer (Thermo Scientific). The mgf files were searched using MeroX to identify crosslinked peptides^69^. The following settings were applied: proteolytic cleavages C-terminally to lysine and arginine; up to 3 missed cleavages; peptide length 5 to 30 amino acids; modifications: alkylation of cysteine by iodoacetamide, oxidation of methionine; crosslinker specificity: lysine, but it can also react with other nucleophilic residues e.g. Tyr, Ser, Thr; precursor precision 10 ppm; fragment precision 20 ppm; signal to noise >2; FDR cut off 1%.

## Data availability

Atomic coordinates have been deposited in the Protein Data Bank, code 9SV8. The cryo-EM of gC2 Δ28-73in complex with C3b has been deposited in the Electron Microscopy Data Bank, codes EMD-55245 (full map) and EMD-55293 (focused classification map).

The mass spectrometry proteomics data have been deposited to the ProteomeXchange Consortium via the PRIDE [1] partner repository with the dataset identifier PXD068845.

Reviewer access details: Log in to the PRIDE website using the following details:

- Project accession: PXD068845

- Token: qs0IyYYVPS0V

Alternatively, reviewer can access the dataset by logging in to the PRIDE website using the following account details:

- Username

- Password: 7wDsyBP10DHN

## Acknowledgements

We thank the Astbury Biostructure Laboratory electron microscopy facility for support with cryo-EM data acquisition, Nasir Khan for support with circular dichroism and John Lambris for reading the manuscript. This project was supported by grant PID2023-149259NB-I00, funded by MICIU/AEI/ 10.13039/501100011033 and by “ERDF A way of making Europe” (to J.F.) and by NIH NIAID grant RO1 AI139618 (to H.M.F.). M.H.R.R. was funded by CONAHCYT (studentship #773992). A. K. C. acknowledges support from an EPSRC studentship (EP/W524372/1 project reference 2879844). A. N. C. acknowledges support of a Sir Henry Dale Fellowship jointly funded by the Wellcome Trust and the Royal Society (220628/Z/20/Z). Electron Microscopy was performed at the Astbury Biostructure Laboratory (University of Leeds), which was funded by the University of Leeds and the Wellcome Trust (108466/Z/15/Z, 090932/Z/09/Z, 221524/Z/20/Z). Funding from Wellcome (223810/Z/21/Z) enabled the purchase of mass spectrometry equipment. Funding from the Wolfson Foundation (PR/jw/md/22597) supported the purchase of the mass photometry equipment.

---

**Extended Data Figure 1.**
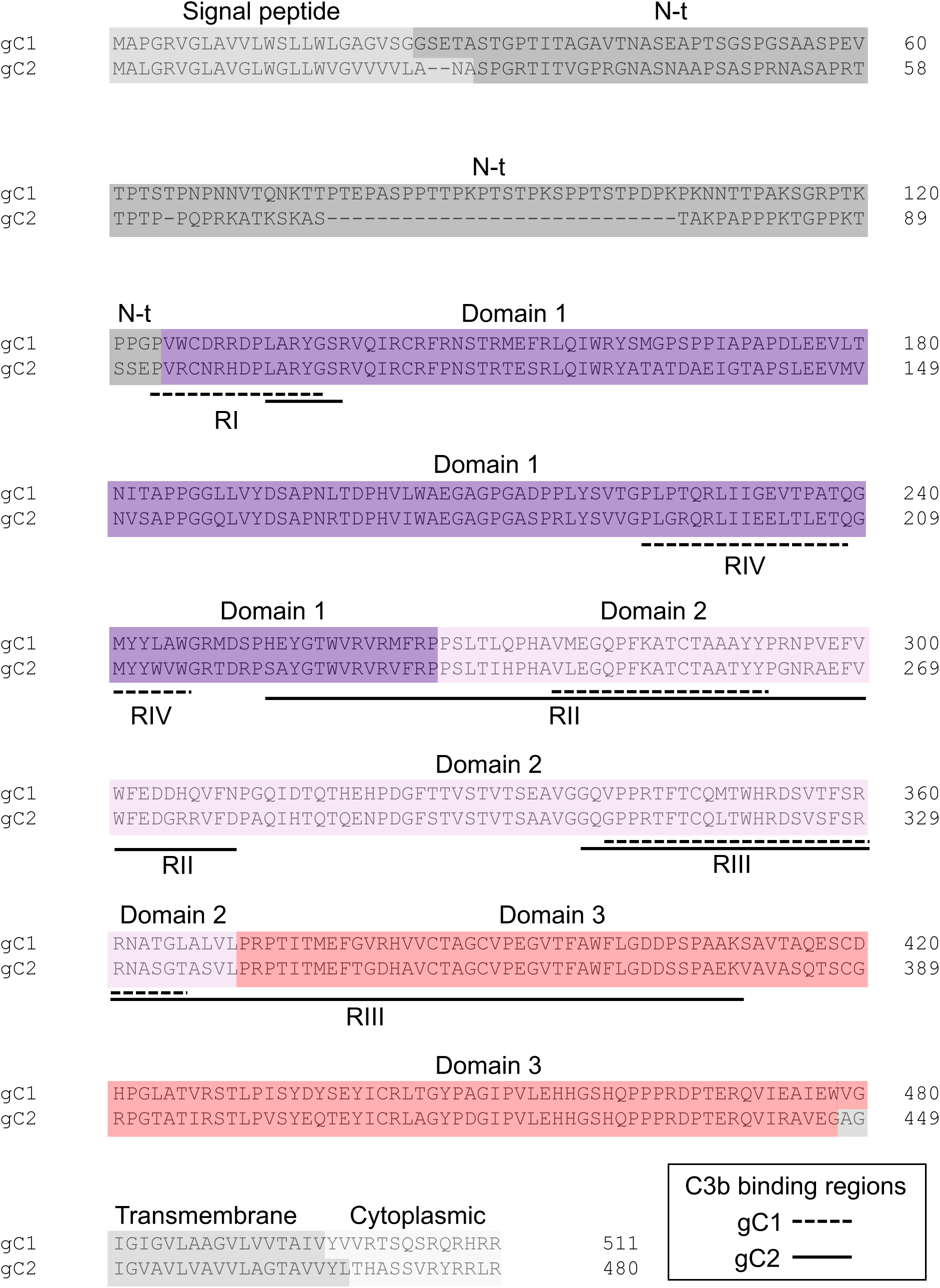
gC1 and gC2 sequence comparison. Sequence alignment of gC1 and gC2, highlighting the different domains (coloured as in Figure 1) and C3b binding regions described in the literature (shown by a dashed line for gC1 and by a continuous line for gC2).

**Extended Data Figure 2.**
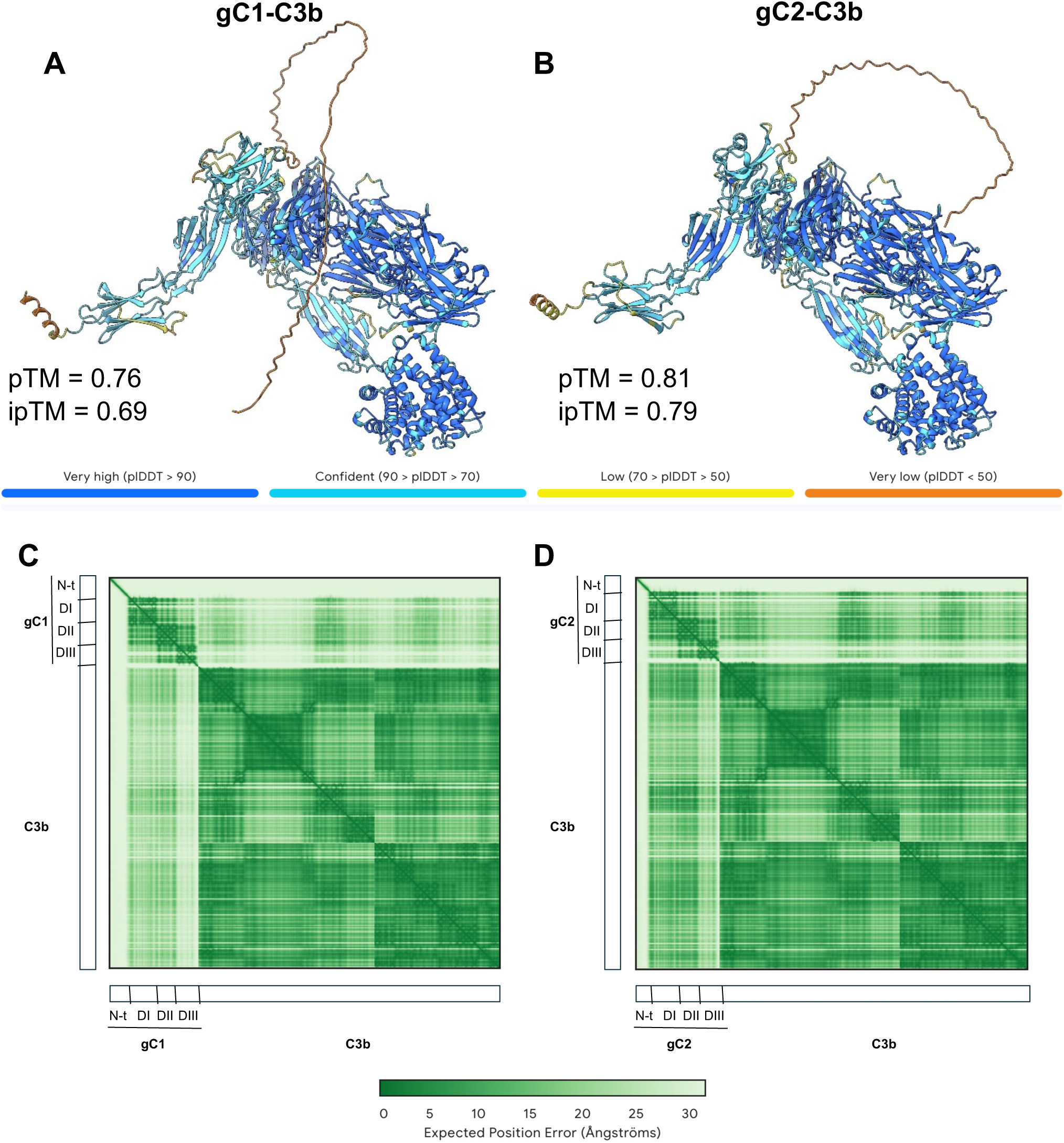
AlphaFold predictions of gC1 and gC2 in complex with C3b. A and B) Comparison of the Alphafold predictions for the gC1-(A) and gC2-(B) C3b complexes, coloured according to pLDDT. pTM and ipTM values for each prediction are shown. C and D) PAE plots for the gC1-(C) and gC2-(D) C3b complexes.

**Extended Data Figure 3.**
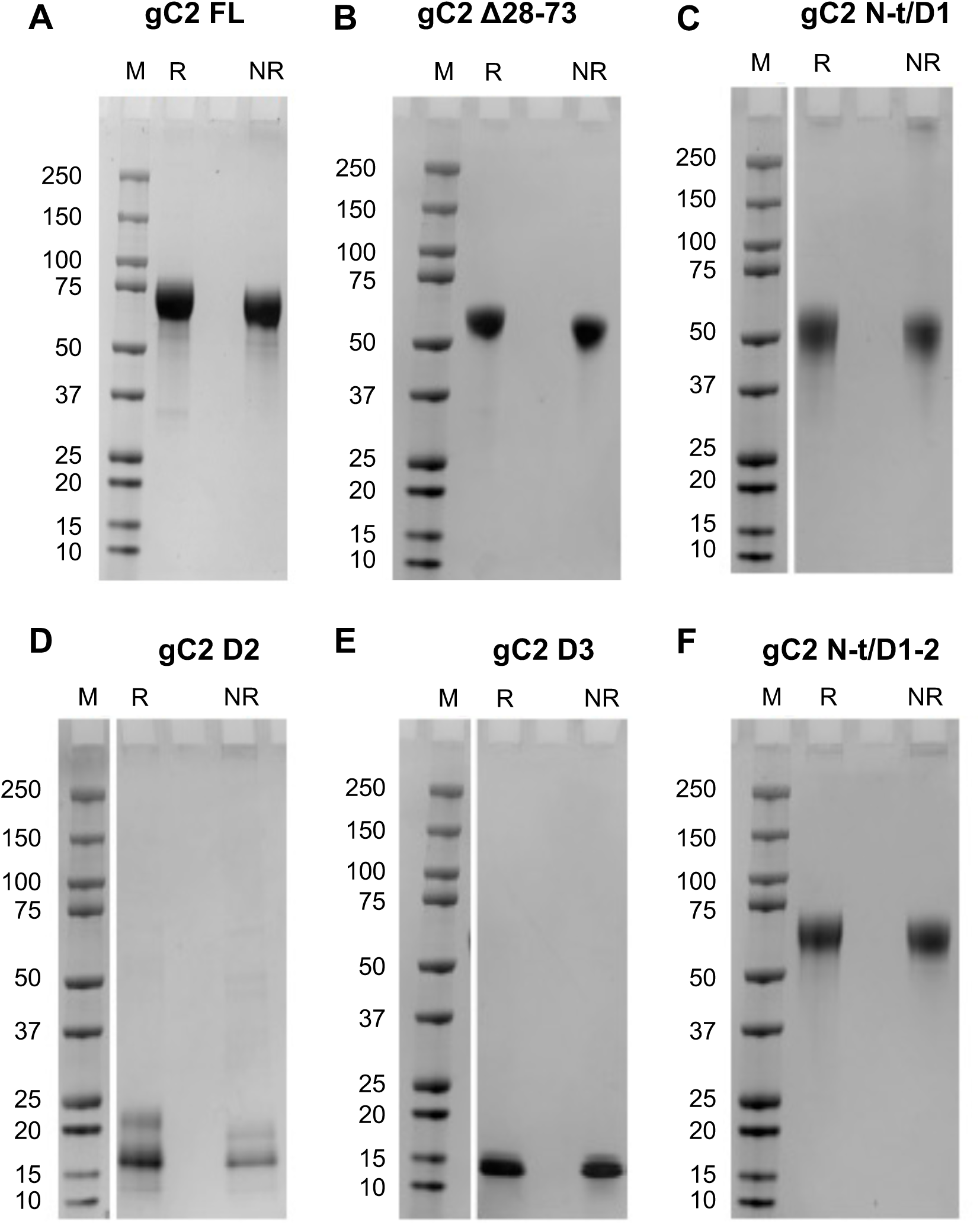
Expression of the gC2 constructs. A-F) SDS-PAGE under reducing (R) and non-reducing (NR) conditions for each gC2 construct. M, molecular weight markers. Proteins were stained using Coomasie Blue.

**Extended Data Figure 4.**
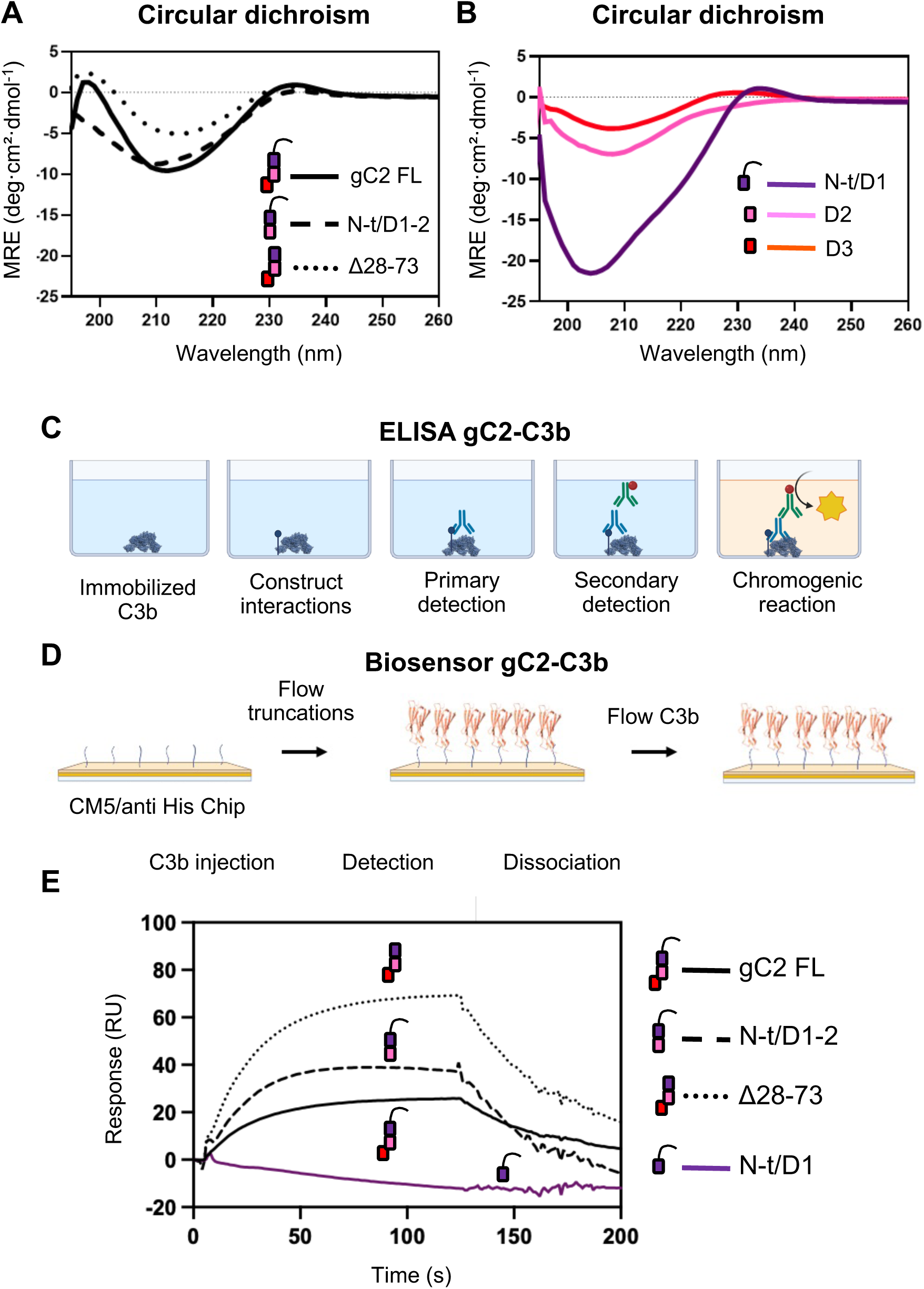
Folding of gC2 constructs and their interaction with C3b. A and B) CD results for the different gC2 constructs. C) Schematic of the ELISA approach used to detect gC2-C3b interactions. D and E) Schematic (D) and results (E) of the biosensor approach used to detect gC2-C3b interactions. Results for each gC2 construct are shown by different lines, as indicated.

**Extended Data Figure 5.**
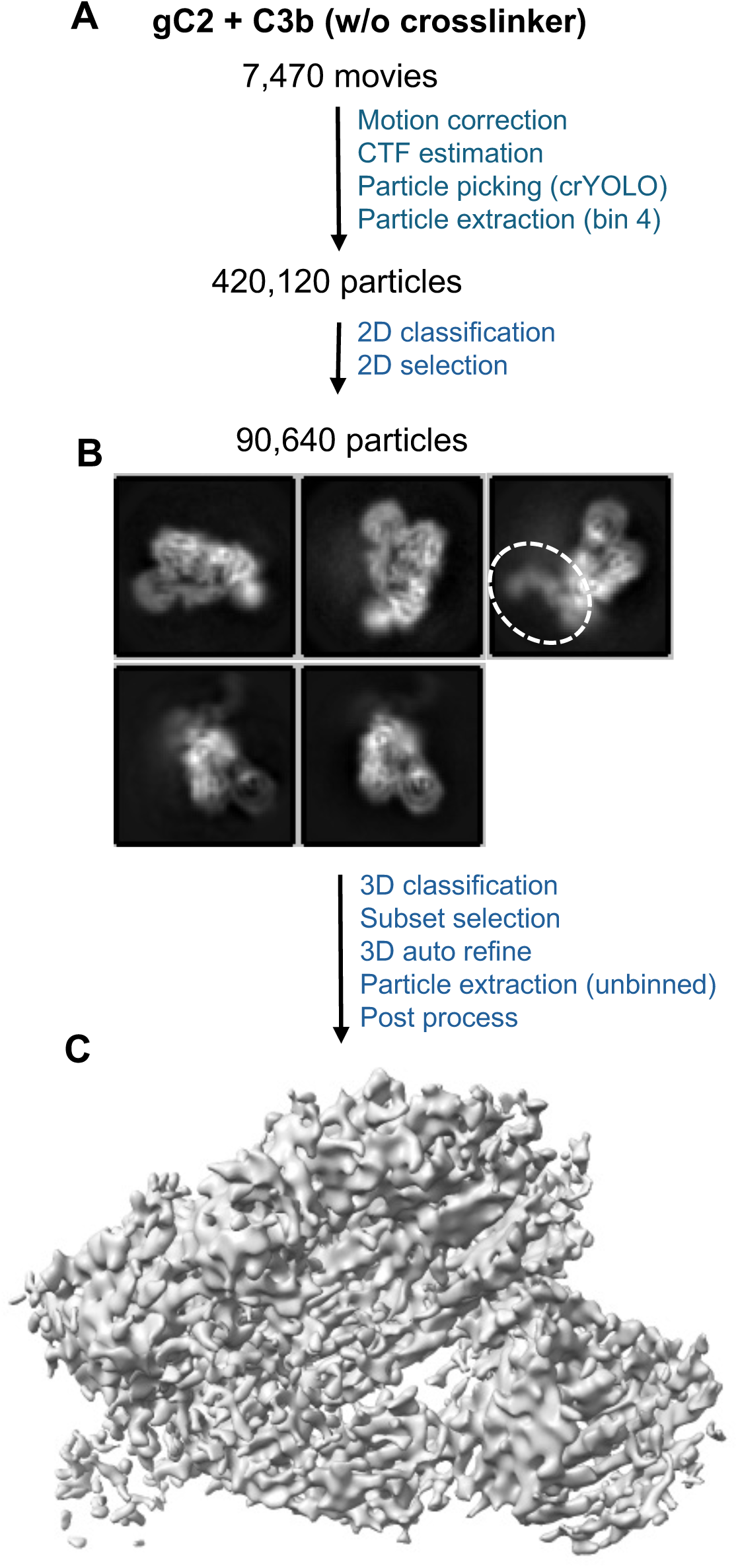
Workflow for cryo-EM data acquisition and image processing for the gC2(426t)-C3b complex in the absence of crosslinker. A) Summary of data acquisition and initial image processing steps. B) Representative 2D classes, highlighting faint densities for gC2(426t). C) 3D map, without densities corresponding to gC2(426t). Image processing was done in Relion.

**Extended Data Figure 6.**
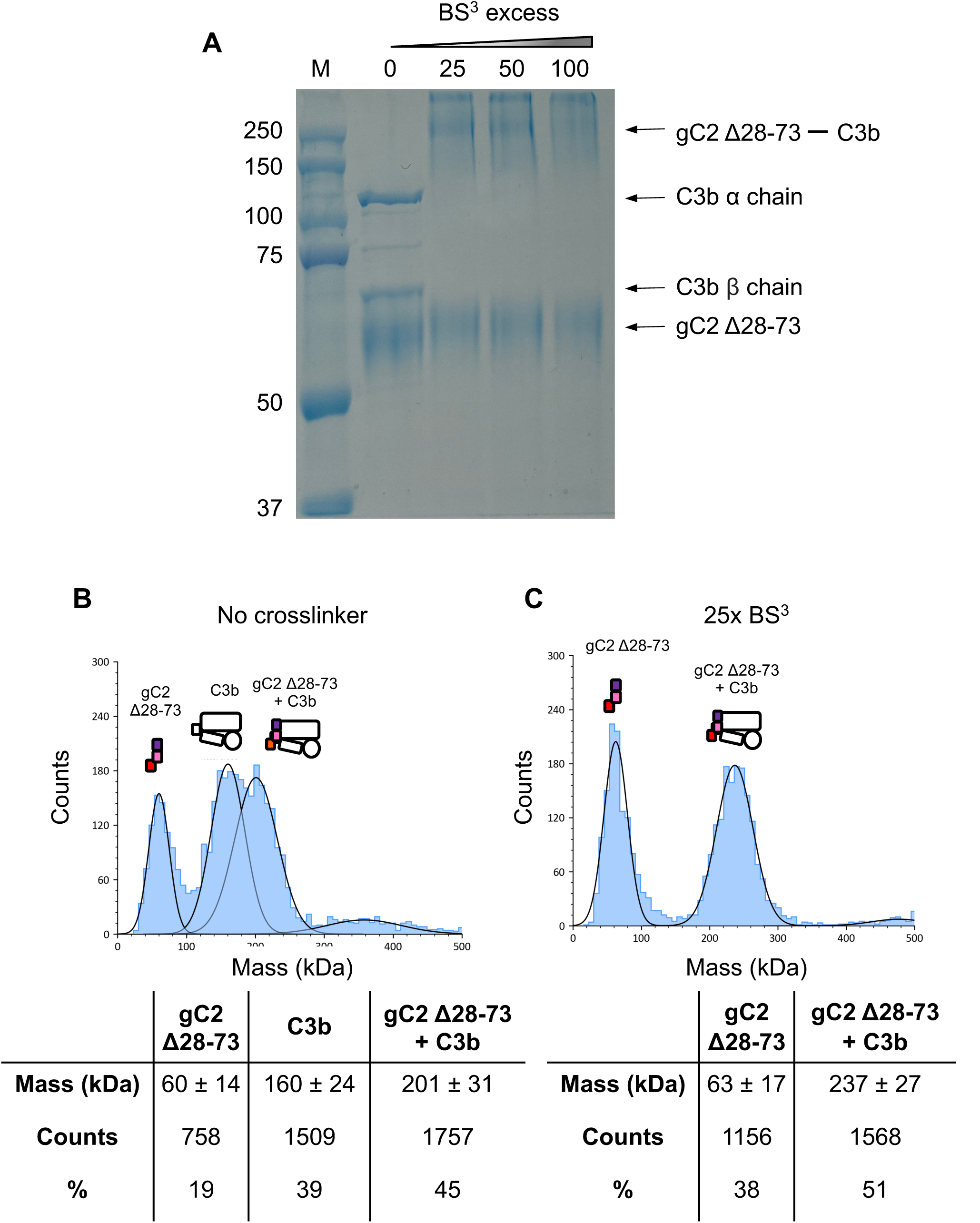
gC2 Δ28-73-C3b crosslinking analysis. A) SDS-PAGE of gC2 Δ28-73-C3b at different molar excess of the BS^3^ crosslinker. M, molecular weight markers. B and C) Mass photometry results of the species present after 30 min of incubation of gC2 Δ28-73-C3b in the absence (B) or presence of 25 molar excess of BS^3^ (C). Statistics of the mass photometry results are shown at the bottom for each condition.

**Extended Data Figure 7.**
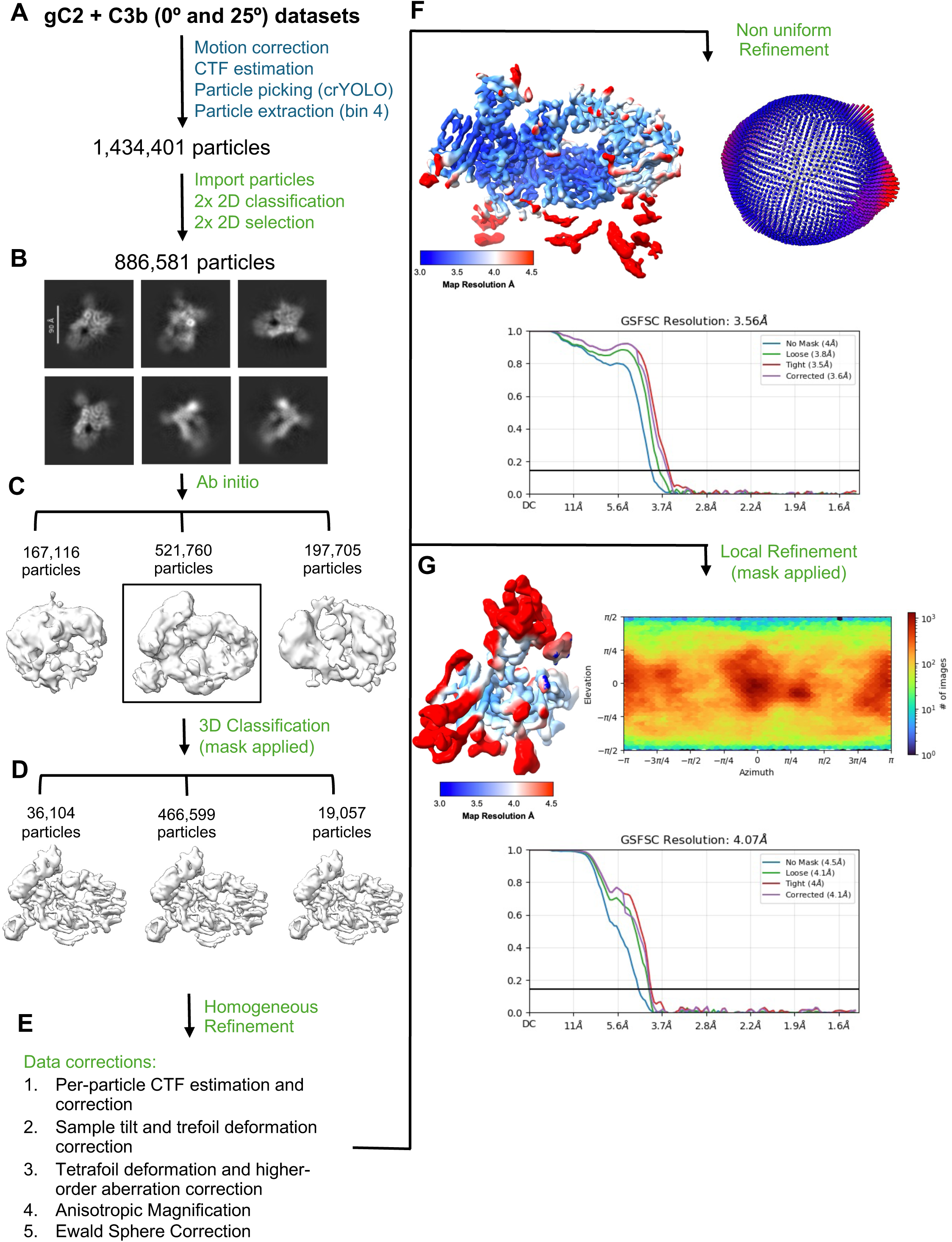
Processing pipeline of the gC2 Δ28-73-C3b cryo-EM data. A) Summary of the initial image processing steps. B) Representative 2D classes. C) Ab initio 3D models. Selected model is highlighted. D) 3D classification. Selected 3D class is highlighted E) Summary of the final image processing steps. F and G) Results for the global (F) and local refined (G) gC2 Δ28-73-C3b maps. Left, final map colour-coded according to local resolution. Right, particle orientation distribution. Bottom, gold-standard FSC curve. Processing jobs were carried out in both RELION v4.0.0 (blue) and cryoSPARC (green).

**Extended Data Table 1.**

| **Data Collection:** | **gC2 Δ28-73 - C3b 0°** | **gC2 Δ28-73 - C3b 25°** |
| --- | --- | --- |
| Microscope | FEI Titan Krios |  |
| Detector | Falcon 4i / Selectris |  |
| Voltage (kV) | 300 |  |
| Nominal magnification | 96k |  |
| Defocus range (μm) | -0.9 to -2.7 |  |
| Pixel size (Å/pixel) | 0.74 |  |
| Electron dose (e<sup>-</sup>/Å<sup>2</sup>) | 43 |  |
| Electron dose per frame (e<sup>-</sup>/Å<sup>2</sup>) | 0.8 |  |
| Exposure time (seconds) | 3.28 |  |
| No. of frames | 20 |  |
| Micrographs collected | 25,000 | 10,087 |
| **Data processing:** |  |  |
| Symmetry | C1 |  |
| Final particle number | 466,599 |  |
| Map average resolution (Å, 0.143 FSC threshold) | 3.56 (4.07 for local refined map) |  |
| Map sharpening Bfactor (Å<sup>2</sup>) | 215.2 (284.7 for local refined map) |  |
| **Atomic model:** |  |  |
| Initial model used | AlphaFold3 (gC2-C3b) |  |
| **Model composition:** |  |  |
| Chains | 3 |  |
| Amino acid residues | 1131 |  |
| Non-hydrogen protein atoms | 8928 |  |
| **Validation:** |  |  |
| Molprobity score | 2.17 |  |
| Clashscore | 7.71 |  |
| Rotamers favoured (%) | 97.5 |  |
| Outliers (%) | 0 |  |
| **Ramachandran plot** |  |  |
| Favoured (%) | 97.5 |  |
| Outliers (%) | 0 |  |
| **Model vs. Map** |  |  |
| FSC (0.5) (Å) | 3.5 |  |
| CC mask | 0.71 |  |

**Extended Data Figure 8.**
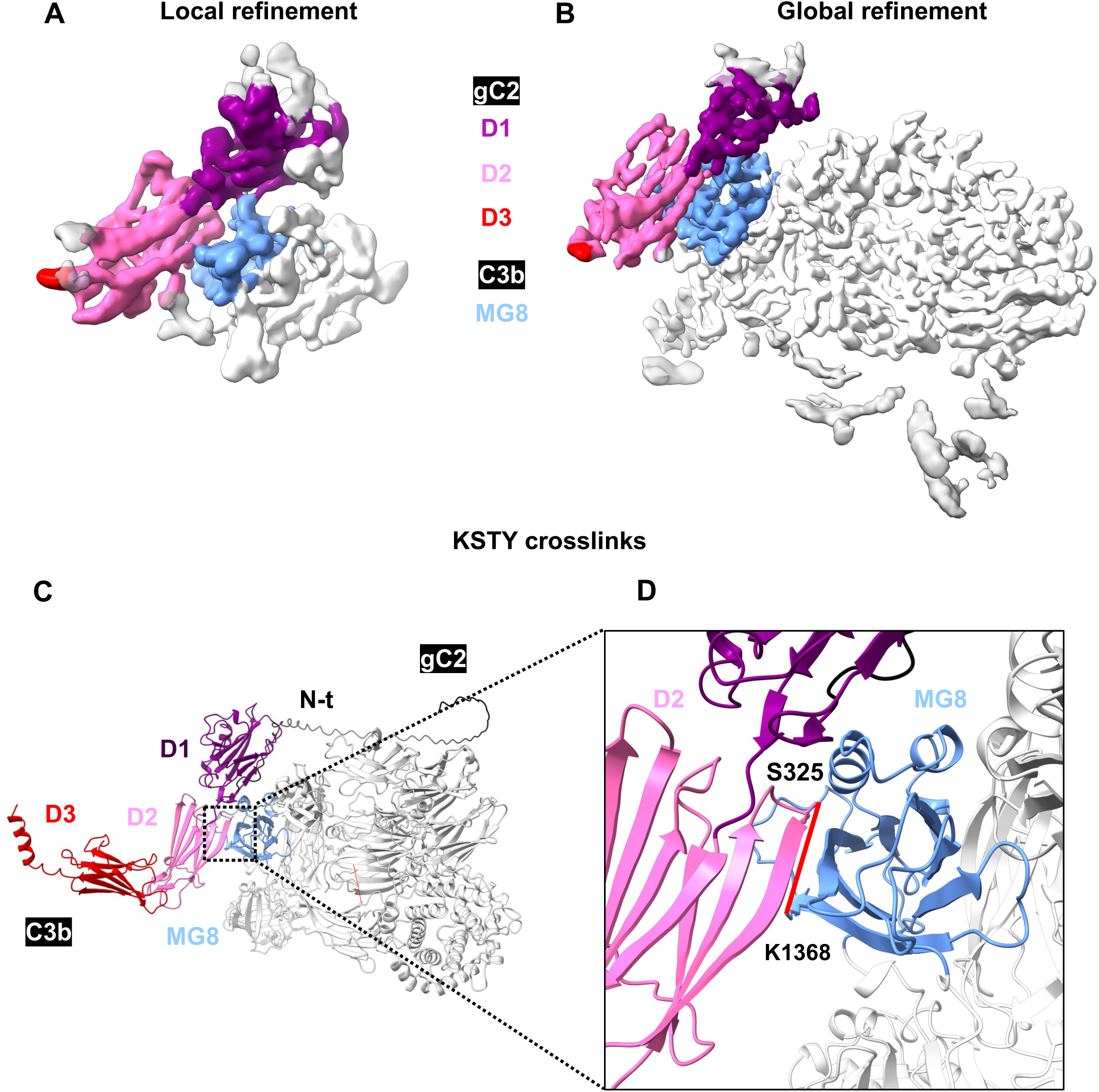
Final cryo-EM maps and additional XL-MS crosslinks. A and B) Final maps for the local (A) and global (B) refinements of the gC2 Δ28-73-C3b complex. Domains are coloured as in Figure 1. C and D) Additional crosslink (red line) between gC2 FL and C3b detected by XL-MS.

**Extended Data Figure 9.**
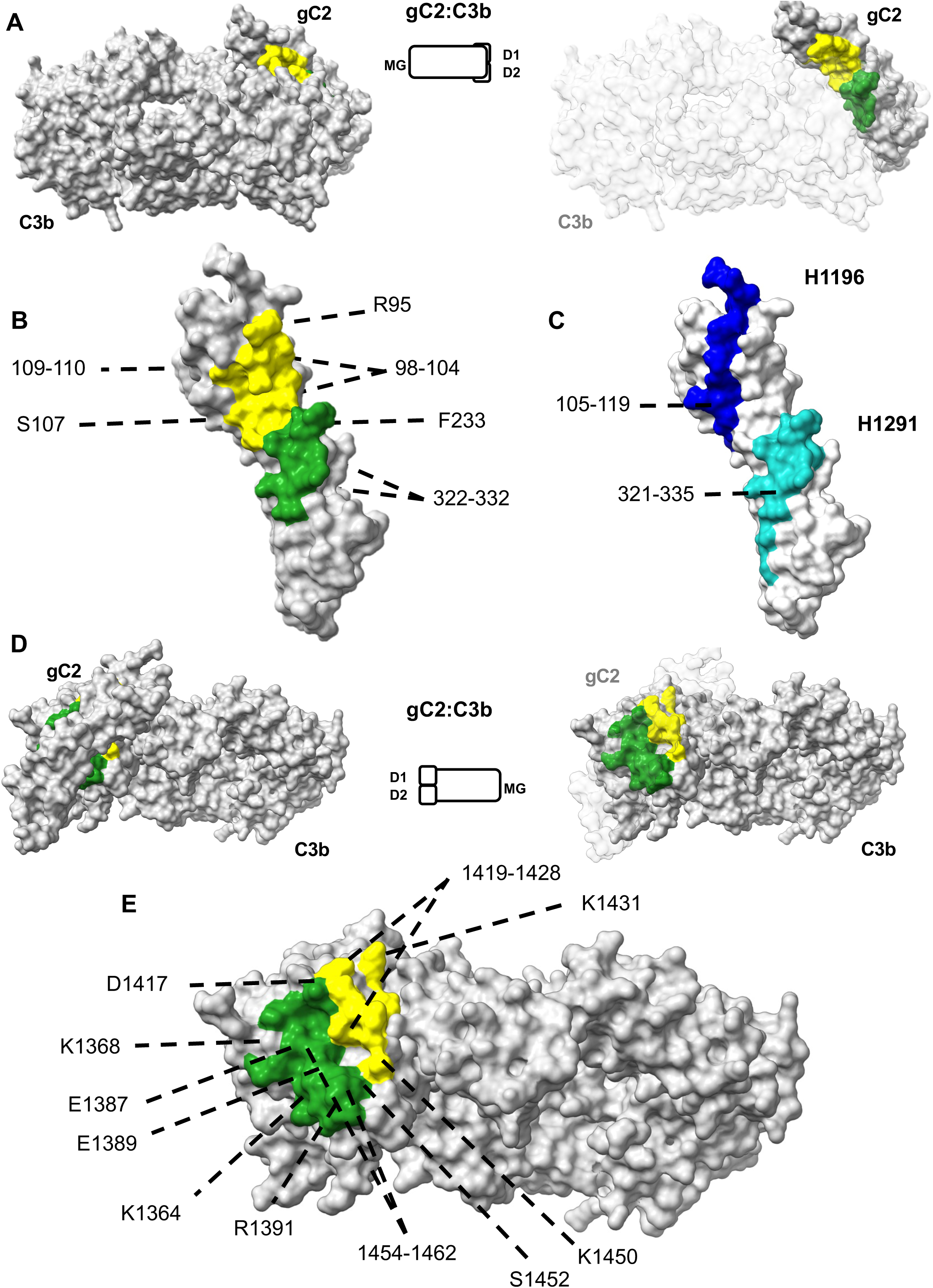
Surface interactions between gC2 and C3b. A and B) Depiction of the gC2 surface that interacts with C3b. gC2 D1 – C3b interactions are shown in yellow, while gC2 D2 – C3b interactions are shown in green. A) Left, complex seen from the C3b side, mostly occluding gC. Centre, schematic representation of the view. Right, complex shown as in the left panel, with transparent C3b. B) Close up view of gC2 as in A, without C3b. C) gC2 linear epitopes recognised by H1196 (dark blue) and H1291 (cyan). To allow a comparison with the C3b-binding region, gC2 is oriented as in B. D and E) Depiction of the C3b surface that interacts with gC2. Interactions are colour-coded as in A. D) Left, complex seen from the gC2 side, partially occluding C3b. Centre, schematic representation of the view. Right, complex shown as in the left panel, with transparent gC2. E) Close up view of C3b as in D, without gC2. In A-B and D-E,

**Extended Data Figure 10.**
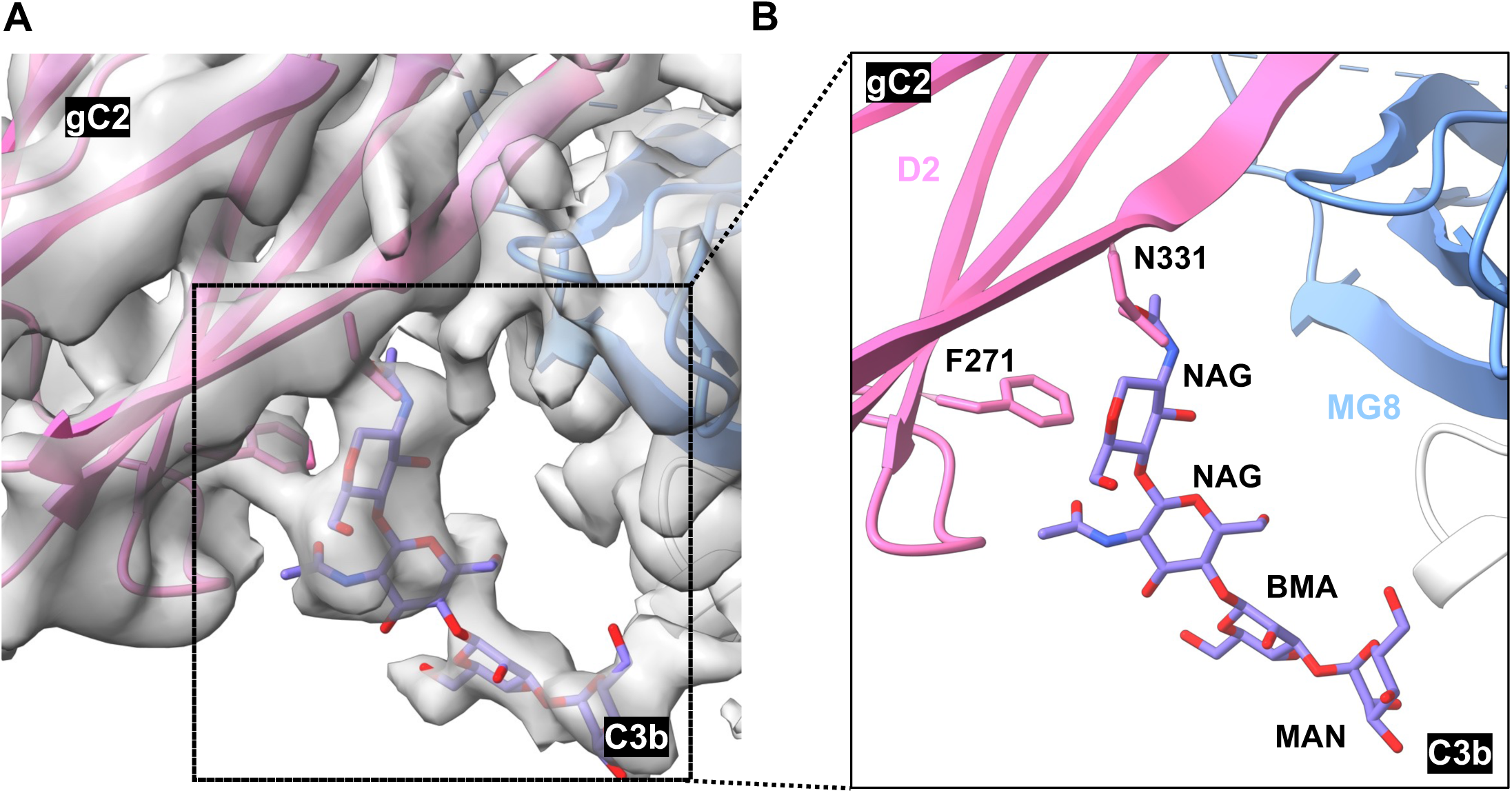
Potential glycosylation of gC2 N331. The extra density around gC2 residue N331 could accommodate at least 4 glycans. The identity of these residues could not be determined due to the resolution of the cryo-EM map in this region. A) Fit of 4 modelled glycans into the cryo-EM map. B) Modelled glycans.

**Extended Data Figure 11.**
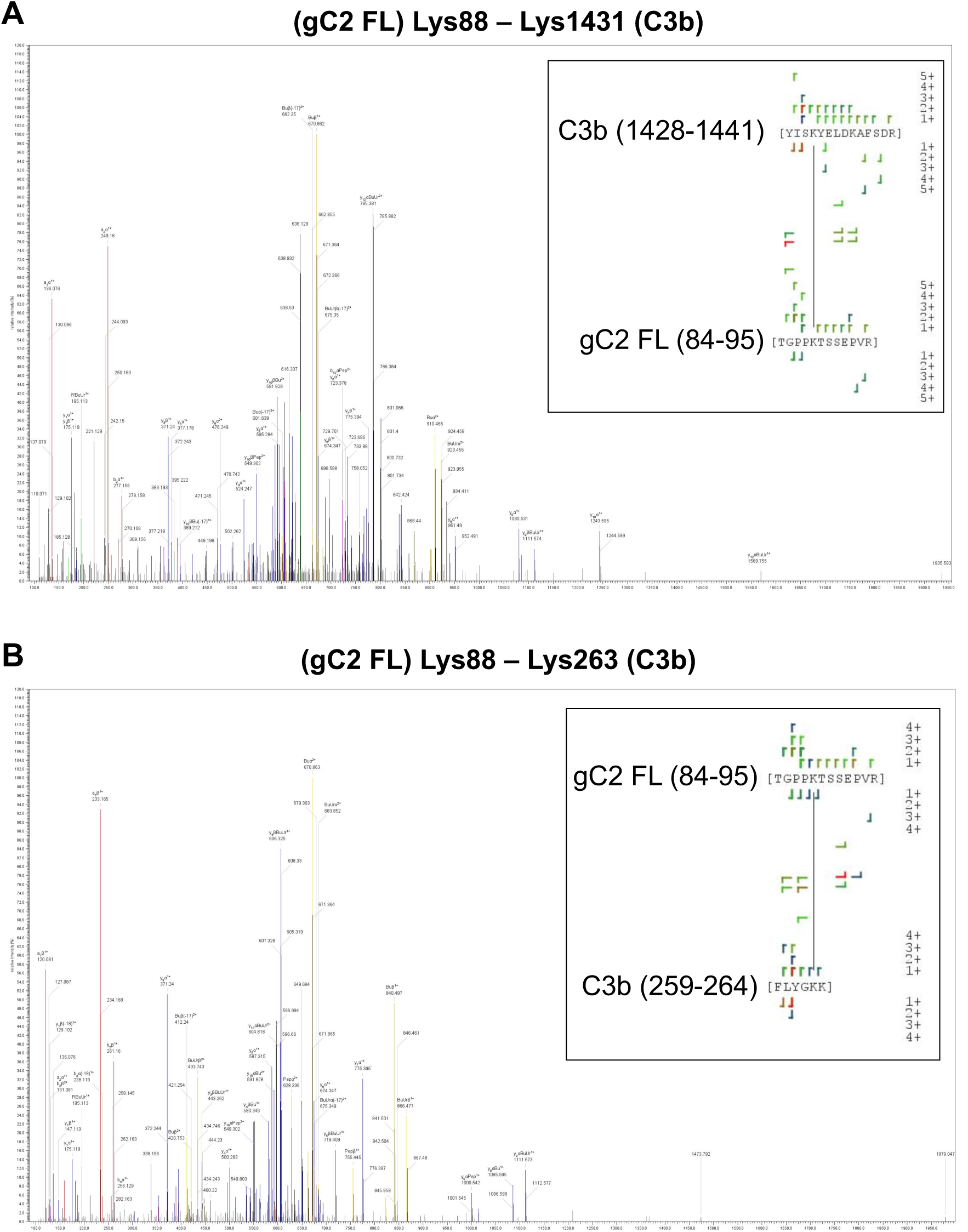
Exemplar MS/MS spectra of identified interprotein crosslinks of the gC2 FL-C3b complex.

**Extended Data Figure 12.**
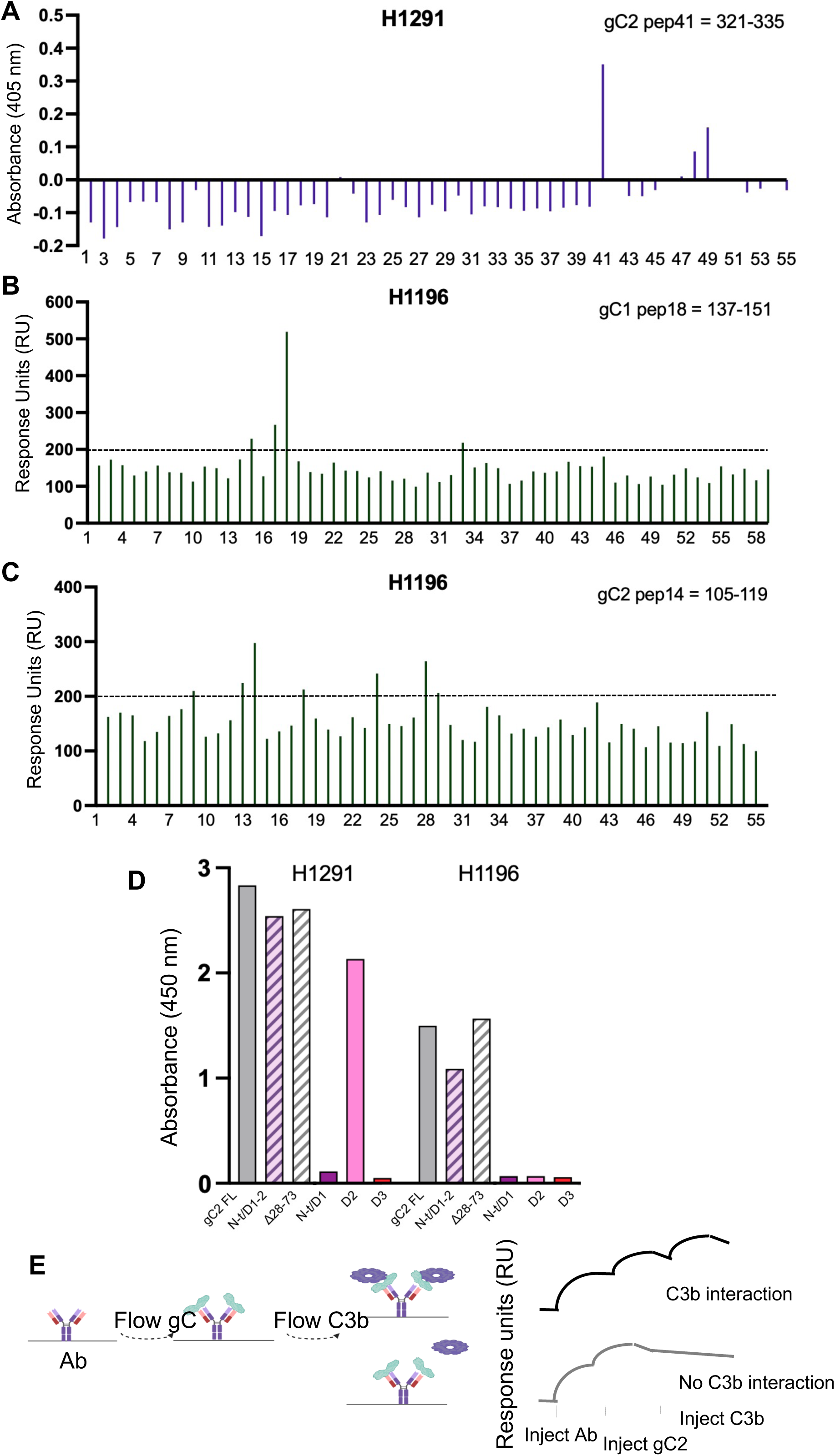
Analysis of epitopes recognised by gC2 antibodies. ELISA results of H1291 (A) and SPR (Caterra LSA) results of H1196 (B and C) screened against 15 amino acid peptides of gC2 (A and C) and gC1 (B). D) ELISA results of H1291 and H1196 when incubated with different gC2 constructs. E) Schematic representation of SPR (Biacore 1k+) experiments to characterise gC2-C3b interaction blocking.

